# Stochastic model for cell population dynamics quantifies homeostasis in colonic crypts and its disruption in early tumorigenesis

**DOI:** 10.1101/2023.03.19.533357

**Authors:** Konstantinos Mamis, Ruibo Zhang, Ivana Bozic

## Abstract

The questions of how healthy colonic crypts maintain their size, and how homeostasis is disrupted by driver mutations, are central to understanding colorectal tumorigenesis. We propose a three-type stochastic branching process, which accounts for stem, transit-amplifying (TA) and fully differentiated (FD) cells, to model the dynamics of cell populations residing in colonic crypts. Our model is simple in its formulation, allowing us to estimate all but one of the model parameters from the literature. Fitting the single remaining parameter, we find that model results agree well with data from healthy human colonic crypts, capturing the considerable variance in population sizes observed experimentally. Importantly, our model predicts a steady state population in healthy colonic crypts for relevant parameter values. We show that *APC* and *KRAS* mutations, the most significant early alterations leading to colorectal cancer, result in increased steady-state populations in mutated crypts, in agreement with experimental results. Finally, our model predicts a simple condition for unbounded growth of cells in a crypt, corresponding to colorectal malignancy. This is predicted to occur when the division rate of TA cells exceeds their differentiation rate, with implications for therapeutic cancer prevention strategies.

## 1 Introduction

The epithelial inner surface of the human colon exhibits about 10^7^ to 10^8^ column-like invaginations [1], called crypts, each one of which contains ∼ 2000 cells [2, 3]. In order to maintain their total size, mature colonic crypts are in homeostatic steady-state, where the cell proliferation in the lower part of the crypt is in equilibrium with the cell loss due to apoptosis [3, 4, 5]. Depending on their proliferation capability, cells in colonic crypts are identified as stem, transit-amplifying (TA), or fully differentiated (FD) [4, 6]. In the base of the crypt column resides a small population of stem cells [1, 7], which can divide and produce cells that are transit amplifying. Cell proliferation in colonic crypts is largely maintained by the TA cells. TA cells divide rapidly as they migrate upward, towards the crypt lumen [8]. After undergoing several rounds of divisions [9], they lose their proliferation capability, resulting in fully differentiated cells [10, 11]. Finally, FD cells undergo apoptosis and are shed into the intestinal lumen [5, 12]. Except for stem cells, epithelial cells reside in the crypt only for a few days, making the intestinal epithelium one of the most rapidly renewing tissues in the human body.

So far, many models have been proposed for the cell population dynamics in a crypt [13], with a lot of works focusing on stem cell dynamics in particular [14]. Cell population models for crypts broadly fall into two categories: spatial and compartmental models. Spatial models are based on the fact that the proliferation capability of each cell diminishes gradually as it moves upwards in the crypt, and they typically take the stochasticity of cell dynamics into account [3, 15, 16, 17, 18]. The main drawback of spatial models is their high level of complexity, since they may include the cell proliferation gradient along crypt’s vertical axis [3, 15], a detailed description of the cell mitotic cycle [16, 17, 18], cell mobility [17] or Wnt signalling [18]; however for the parameters of these processes, estimation from experimental measurements is not straightforward. Thus, spatial models are particularly useful in deducing qualitative results or testing different hypotheses concerning the mechanisms of cell dynamics, rather than making quantitative predictions for the cell populations in a crypt.

Compartmental models, on the other hand, do not make use of the spatial character of cell proliferation, describing instead the progression of cells through the different compartments that constitute the total cell population in a crypt. Most of compartmental models are deterministic [4, 19, 20, 21], and thus cannot capture the variability in cell populations within the crypt, which is significant [2, 3, 15, 22]. Compartmental models have been mainly employed to describe the mechanisms in cell population dynamics that result in homeostasis, their stability, and possible ways they can be disrupted resulting in tumorigenesis [2, 4, 23]. Of particular interest are feedback control mechanisms between different populations of cells [2, 4, 21]. However, most of the results are qualitative, since the values of model parameters remain unspecified [4, 21, 23]. More detailed compartmental models may contain more compartments corresponding to different phases of cell cycle [19, 24], or incorporate both stochasticity and control networks between the different cell compartments [2]. However, most of their parameters are not estimated directly from experimental measurements; rather, parameter values are adjusted so that model simulations are able to reproduce experimental data, such as the distribution of proliferative cells along the crypt axis, or the averages and variances of cell populations in a crypt.

In the present work, we propose the branching process depicted schematically in Fig. 1, as a new stochastic model for cell dynamics in a colonic crypt that **i)** is minimal and simple in its formulation, which allows us to directly estimate all model parameters except for one, **ii)** is able to model the cell population dynamics in both healthy and mutated colonic crypts, **iii)** its solution is obtained analytically. More specifically, for healthy crypts, our model explains both the attainment of homeostatic steady-state and the variability in cell population sizes from crypt to crypt. Furthermore, our model describes the increase in size of mutant crypts [22], by considering the effects on cell properties of the most common mutations found in early stages of colorectal tumorigenesis, such as *APC* and *KRAS* [25, 26, 27]. Its analytic solution is another significant aspect of the model, as it provides a simple mathematical description of the homeostatic mechanisms inside a colonic crypt and the disruption thereof, shedding light on the mechanisms that result in colorectal tumorigenesis.

**Figure 1:**
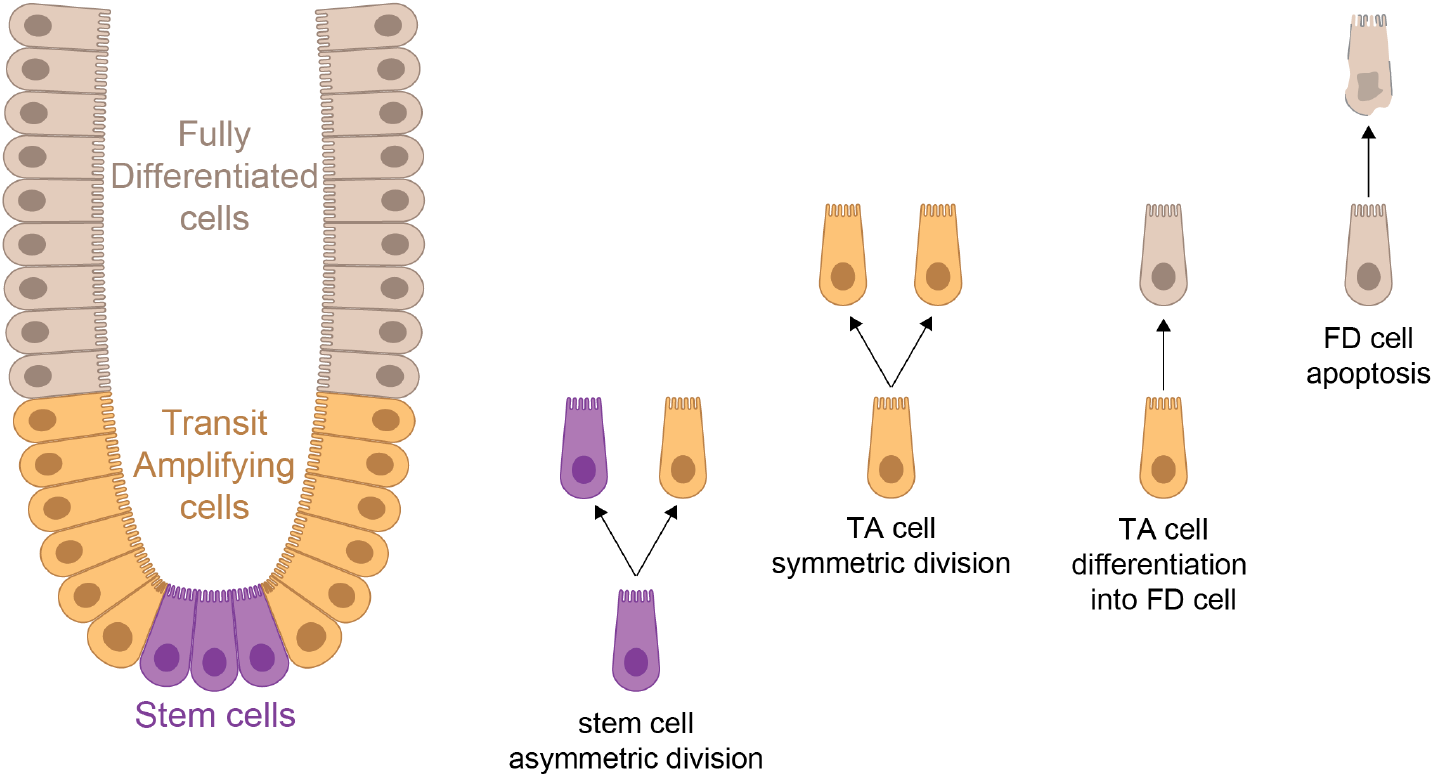
Schematic representation of cell populations and cell dynamics in a colonic crypt.

Branching processes have been used to model cell population dynamics [28], with applications in the study of mutation acquisition in cancer progression [29, 30, 31, 32], tumor progression and ctDNA shedding [33], and epidermal homeostasis [34]. In particular, epidermal homeostasis was successfully described by using an exactly solvable, two-type branching process for cell dynamics [34, 35]. Our model is another example of describing a homeostatic mechanism in healthy tissue by a branching process. From its solution, we are able to determine the steady-state distributions of different cell population sizes in a crypt, as well as exact formulas for their averages and variances.

For the model parameters of stem cell number, stem and TA cell division rates and FD cell apoptosis rate, we use typical values found in the literature on healthy human colonic crypts. Since no such direct value estimation was found for TA differentiation rate into FD cells, we estimate it from the average crypt size observed in humans. While we estimate only one parameter using the average crypt size, our model predicts distributions for TA, FD, and total cell populations that are in good agreement with experimental data [2, 3], encapsulating the stochasticity of cell dynamics in a crypt. We also consider the changes in values of model parameters, resulting from the *APC* and *KRAS* mutations that are commonly found in early stages of colorectal tumorigenesis. We show that our model is able to capture the changes in size and population structure of mutant crypts demonstrated experimentally.

## 2 Model

We propose a continuous-time branching process with three types, corresponding to stem (S), transit-amplifying (TA) or fully differentiated (FD) cells, to model population dynamics in a colonic crypt. In this stochastic model depicted in Fig. 1, cells undergo divisions, differentiations and apoptoses with constant rates but at random times. We describe this branching process by the rate equations

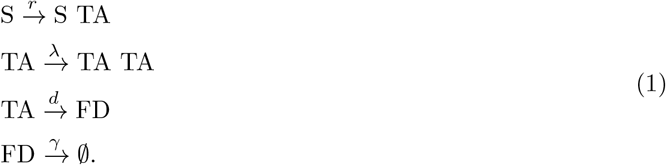

In the model, the number *N*_0_ of stem cells per crypt remains constant, since we consider that stem cells only divide asymmetrically with rate *r*, producing one stem and one TA cell. Recent research [1, 36] has challenged the previously held position that stem cells divide predominantly asymmetrically. Each stem cell is thought to divide symmetrically into two daughter stem cells most of the time, with asymmetry being controlled on the population level: Stemness is a property of cells residing in the stem cell niche in colonic crypt base [37, 10], and each symmetric division results in one surplus stem cell being pushed upwards, differentiating into a TA cell [38, 39]. This mechanism results in the tight regulation of stem cell numbers per crypt, which in turn allows us to assume, in our model, that stem cells *effectively* undergo asymmetric divisions, as each stem cell division ultimately results in one new TA cell.

The proliferating compartment of the crypt consists mainly of the TA cell population. TA cells are rapidly-cycling, undergoing a number of cell divisions before their final differentiation [6]. In our model, TA cell behavior consists of symmetric division with rate *λ*, or differentiation into an FD cell with rate *d*. Note that cell proliferation capability depends on the location of the cell along the crypt axis, since it is controlled by signaling factors whose concentration decreases as cells move away from the crypt base [1]. Thus, in our model we assume that TA differentiations occur independently of TA divisions [40].

Last, FD cells, having lost their proliferation capability, can only undergo apoptosis with rate *γ*.

### 2.1 Average sizes of cell populations and condition for homeostasis

We formulate a system of ordinary differential equations that describes the time evolution of average population sizes of TA and FD cells, denoted as *m*_TA_(*t*) and *m*_FD_(*t*) respectively:

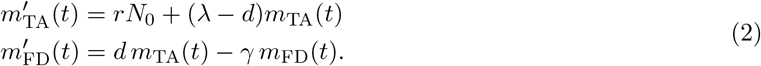

Under condition

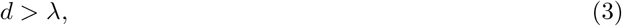

and regardless of the initial cell populations, system of equations (2) attains a steady state; that is, for large times that exceed the cell turnover timescale, the average TA and FD cell populations remain constant and equal to

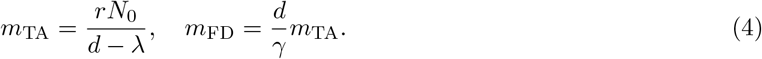

Thus, under condition (3), the crypt maintains a finite size, which, on average, is equal to

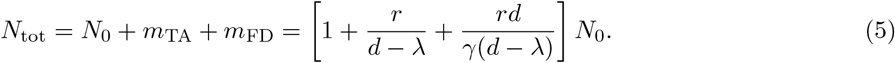

On the other hand, if condition (3) is violated, system (2) predicts an unbounded growth of both TA and FD cell numbers. Thus, stationarity condition *d > λ* provides us with a simple description of the homeostasis for crypt size: In order for the crypt to maintain its finite size, TA cell division has to be regulated, with TA division rate *λ* being less than rate *d* of TA differentiation into FD cells. While we derived the stationarity condition for the averaged Eqs. (2), it is also the stationarity condition for the stochastic population model (1).

### 2.2 Probability distribution for number of TA and FD cells

Moving now to the stochastic setting of model (1), we explicitly determine the exact steady-state probability distribution for the number of TA cells per crypt: For large times, the number of TA cells follows a generalized negative binomial distribution, with the number of successes being *rN*_0_*/λ* and success probability (*d − λ*)*/d* (Supplemental Material, section S3). In other words, probability that the number of TA cells is equal to *n*_1_ is given by

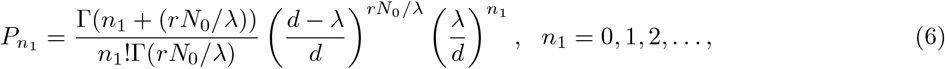

where Γ(·) is the gamma function. The average number of TA cells, as calculated from distribution (6), agrees with the result (4) we obtained from the averaged deterministic problem, while the variance of TA cell population reads

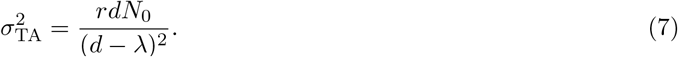

Note that, Dewanji et al. [41] have identified negative binomial distribution as the solution to a similar cell population problem, albeit for normal/mutant bacteria in a generalized Luria-Delbrück model.

Furthermore, by making use of FD cell dynamics, in combination with the known distribution 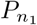 for the TA cell population, we formulate the following approximation for the steady-state distribution of the number *n*_2_ of FD cells (see Supplemental Material, section S4):

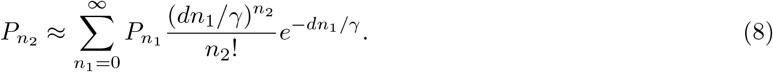

Approximation (8) is easily computable, and we show its accuracy in Fig. 3, for model parameters corresponding to healthy human colonic crypts.

### 2.3 Probability distribution for total number of cells in a crypt

In order to determine the steady-state distribution of the total number of cells per crypt, we formulate and solve the Kolmogorov equation corresponding to branching process (1) (Supplemental Material, section S5). From its solution, we derive the exact distributions for TA, FD and total cell populations in a crypt (Supplemental Material, section S6). The exact distributions for cell populations agree with the results obtained from direct Gillespie simulations of the model (Supplemental Material, Fig. S1).

From the exact solution, we derive closed form formulas for the variance for the number of FD cells

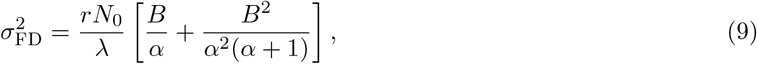

where *α* = (*d − λ*)*/γ, B* = *λd/γ*^2^, and the covariance between the TA and FD cell populations

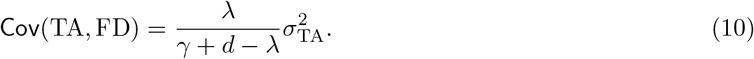

From these results and the variance of TA cell population, we calculate the variance of the total cell population in a crypt:

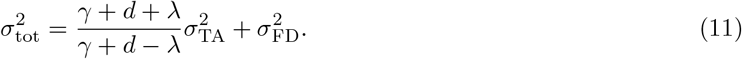

### 2.4 Maintaining homeostasis under different apoptosis rates

Apoptosis rate of FD cells *γ* is the parameter with the greatest uncertainty in the model; estimations for FD cell lifespan may vary between 1 to 6 days [42]. Our model is able to efficiently attain homeostasis (i.e. maintain a constant crypt size distribution) for different FD apoptosis rates *γ*, by adjusting TA differentiation rate *d*. To show this, we consider different values for *γ*, and for each one of these we adjust *d* so that the average crypt size stays ∼2400 cells. We see in Fig. 2 that this adjustment in *d* results not only in keeping the average crypt sizes constant, but also leaves the crypt size distribution effectively unaltered. This is achieved by the simultaneous adjustment of the distributions for TA and FD cells: An increase in apoptosis rate *γ* results in an increase in the average size and variance of TA cell population and a simultaneous decrease in the FD cell population, while a decrease in *γ* has the opposite results. These changes in TA and FD cell populations are in agreement with the remark by Potten and Loeffler [43] that a large TA cell population is needed when cell turnover is high, i.e. for increased apoptosis rates *γ*; if *γ* is low, the tissue could in principle be maintained with just stem and FD cells.

**Figure 2:**
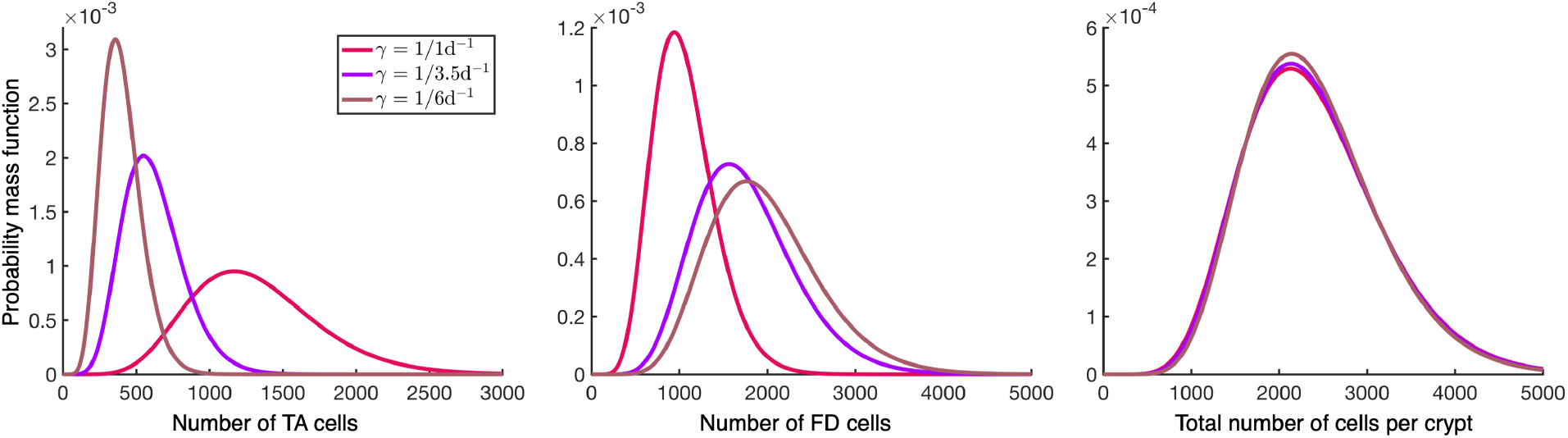
Crypts can maintain homeostatic size distribution under varying apoptosis rates. Steady-state probability distributions of TA, FD, and total cell populations in colonic crypts, for different apoptosis rates *γ*, determined by the solution of branching process (1). Values for parameters *N*_0_, *r, λ* are described in Section 3, and *d* is adjusted to keep the average crypt size constant.

**Figure 3:**
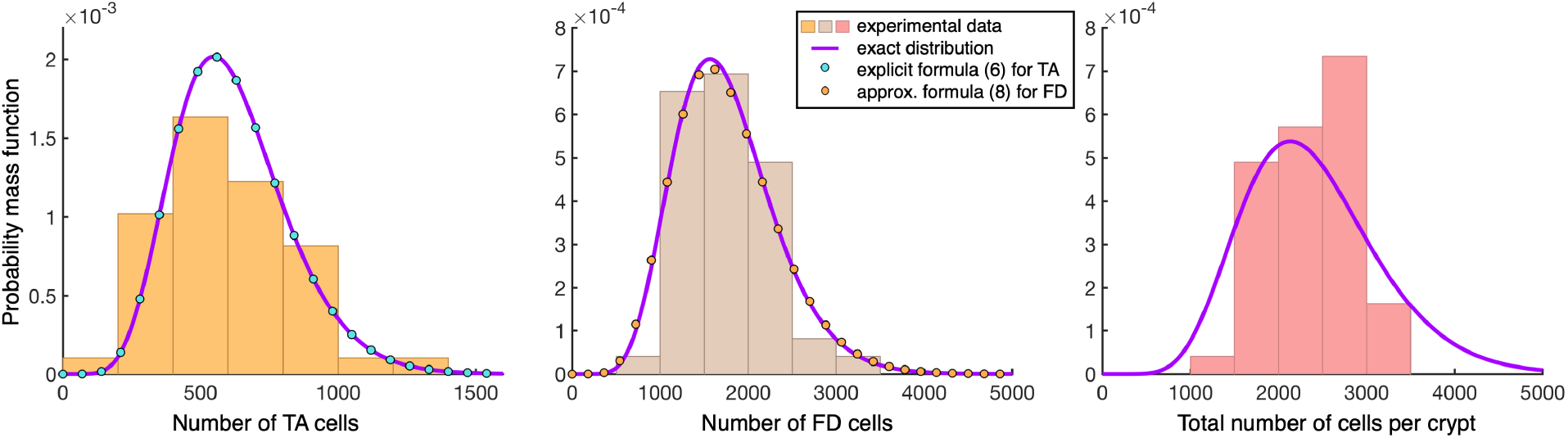
Comparison between experimental data and model results. Steady-state probability distributions of TA, FD, and total cell populations in healthy human colonic crypts. Histograms represent experimental data [3]. Purple curves are the exact distributions determined by the solution of branching process (1), for parameters values that correspond to healthy human colonic crypts: *N*_0_ = 18 cells, *r* = 1*/*2.5d^*−*1^, *λ* = 1*/*30h^*−*1^, *γ* = 1*/*3.5d^*−*1^; *d* is adjusted so that *N*_tot_ ∼ 2400 cells. The explicit formula (6), and the approximate formula (8), for TA and FD cell populations respectively, are denoted by circles in the first and second panels.

## 3 Comparison of model results to experimental data

### Parameter values

For the comparison to data, we calibrate model (1) with parameter values from the literature that are typical to human colonic crypts. Unfortunately, model parameters have been determined more precisely for murine and not human colonic crypts, since most of the experimental studies have been performed on mice. In murine colonic crypts, by identifying that *Lgr5* ^+^ crypt base columnar cells fulfill the stemness criteria, stem cell number has been experimentally determined to ∼15 [38]. For human crypts, the stem cell number estimations are similar or slightly higher than the ones reported for mice, i.e. between 15 and 20 cells [44, 37, 2, 9]. For example, the modal value of stem cells in human crypts is estimated between 15 and 20 stem cells by methylation pattern analysis [9], and an average of 18.7 stem cells has been reported from human biopsy specimens by using biomarkers that stain the active stem cells in a crypt [2]. Thus, for our model, we use the value *N*_0_ = 18. Stem cell division rate *r* has been determined to once a day for mice [10, 8], possibly under the influence of circadian factors [45]. In humans, colonic stem cells divide once every 2-3 days [27, 46]; this estimation is determined from an in vivo bromodeoxyuridine labeling of cells in human colonic crypts [47, 48]. Thus, we consider the value *r* = 1*/*2.5d^*−*1^. In mice, it has been determined that TA cells divide every 12 to 16 hours [45, 6, 25], taking approximately half as long for each of their cycles compared to stem cells [8]. For the human colon, the cell cycle in the mid-crypt section, where the proliferative cells reside, is estimated to 30 hours, via the experimental determination of the labeling index along the crypt axis [47, 48, 49]. From this, we use *λ* = 1*/*30h^*−*1^ for the division rate of TA cells. In the same studies [47, 48, 49], the mean turnover of FD cells has been determined to 3.5 days, with a estimated variability from 2.8 to 5.6 days. In a more recent meta-analysis [42], estimates typically fall in the 1-6 days range for colorectal differentiated cells. Additionally, from an estimated renewal rate of about 100 times per year for the FD population in the colonic crypt, the FD lifespan has been inferred to ∼3-4 days [50]. Thus, we consider *γ* = 1*/*3.5d^*−*1^ as the apoptosis rate for FD cells.

Last, we found no direct estimation in the literature for the rate *d* of TA cells differentiation into FD cells. We treat *d* as a fitting parameter for our model, so that the average total size *N*_tot_ of the crypt is ∼2400 cells, matching experimental observations [2, 3]. This is performed by using Eq. (5) for average crypt size, resulting in the value *d* = 0.0338h^*−*1^. A similar fitting has been performed for a deterministic compartmental model for cell dynamics in hematopoiesis [20].

We compare our model predictions with the data from 49 colorectal crypts obtained from human biopsy specimens [2, 3]. In Table 1, we compare results for average sizes and variances of TA, FD, and total cells per crypt, to the corresponding values from the experimental data set. In Fig. 3, we show the model results for probability distributions of cell populations, along with the histograms of the experimental data, and we observe that they are in good agreement. More specifically, by performing the chi-square goodness of fit test at 5% significance level (Supplemental Material, section S7), we find that the data are consistent with the distributions predicted by our model. The agreement between the theoretical predictions of the model and the size distributions of the three cell populations (TA, FD and total number of cells) is all the more striking if we recall that it relies on fitting only a single parameter.

**Table 1:** Comparison of sizes of cell populations in healthy human colonic crypts from model and experimental data.

| Cell type | Average | Variance | CV%* |
| --- | --- | --- | --- |
| TA cells (data) | 608.9 | $5.5 \cdot 10^4$ | 38.4 |
| TA cells (model) | 618.1 | $4.3 \cdot 10^4$ | 33.6 |
| FD cells (data) | 1768.2 | $1.9 \cdot 10^5$ | 24.6 |
| FD cells (model) | 1756.0 | $3.3 \cdot 10^5$ | 32.8 |
| total number of cells (data) | 2392.1 | $2.4 \cdot 10^5$ | 20.5 |
| total number of cells (model) | 2392.1 | $6.0 \cdot 10^5$ | 32.5 |

\*Coefficient of variation (CV%) is the ratio of the standard deviation to average, expressed as percentage.

The main discrepancy between our model results and data from [3] is the correlation between the TA and FD cell populations. Pearson’s correlation coefficient, as calculated from data, is 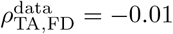, while the model predicts 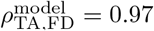. The observation that numbers of TA and FD cells are uncorrelated has been attributed to a feedback mechanism in cell population dynamics [2]. In our model, the correlation between TA and FD cell populations can be reduced significantly if we allow apoptosis rate *γ* to vary randomly from crypt to crypt. More specifically, by considering FD cell lifespan in each crypt to follow a continuous uniform distribution between 1 and 6 days (thus having an average of 3.5 days), and keeping the average crypt size constant, correlation coefficient is reduced to *ρ*_TA,FD_ = 0.28, while the cell population distributions stay close to the experimental data. Moreover, the distributions for FD and total numbers of cells, as predicted by the model for constant and varying *γ*, effectively coincide (see Supplemental Material, Fig. S2).

On the other hand, the proportional growth between total crypt size and the proliferative cell compartment has been experimentally observed [51], and has also been incorporated in spatial crypt models [15]. This experimental result is in agreement with the positive linear dependence between TA and total cell population in our model. The reason for such potentially conflicting experimental results could potentially be the identification of proliferating cells as the cells that are Ki67 positively staining [3]; Ki67 staining is not a perfect differentiator between TA and FD cells, since some TA cells may not be stained due to the cell-cycle phase they currently are, and thus counted as FD cells [52].

### 3.1 Insights into the number of stem cells in crypt

Before turning to study the effects of driver mutations, we discuss the implications of our model on the potential numbers of stem cells in healthy human colonic crypts. We note that there is an ambiguity in the field regarding the number of stem cells per crypt, since lower values for *N*_0_ around 5 have been reported [53, 54, 27]. We employed our model to investigate if it can provide insight into the likely stem cells number in a crypt. For *N*_0_ = 5, our model predicts distributions for cell populations that are not in agreement with experimental data (Supplemental Material, section S7 and Fig. S3); chi square goodness of fit test at a 5% significance level disproves the hypothesis that experimental data for FD and total cell populations are consistent with the distributions from model results with *N*_0_ = 5. Importantly, for *N*_0_ = 5, our model predicts coefficients of variation (CV) for all cell populations ∼ 63% This is significantly higher than the values of CV calculated from experimental data (Table 1). Thus, our model results corroborate the experimental findings that there are ∼ 18 stem cells per crypt [3, 2].

## 4 Effects of driver mutations on population dynamics

Having described the steady-state of healthy colonic crypts, we now use model (1) to determine the size and composition of mutated crypts in early stages of colorectal tumorigenesis, i.e. after the acquisition of certain genetic driver mutations such as *APC* and *KRAS. APC* inactivating mutation is sufficient for the initiation of early colorectal adenomas [55], and is present in ∼ 80% of human colorectal cancers [56], resulting in the increase in proliferating cell compartment and the total crypt size [57, 58]. *KRAS* activation [59] is present in ∼ 40% of human colorectal cancers and large adenomas [60], while it is often detected in human hyperplastic polyps [61]. Despite the structural alterations, mutated crypts retain the cell hierarchy described for healthy crypts [6, 25, 27].

The main effect of *Kras* mutation is a 60% increase in stem cell division rate [25], while the number of stem cells per crypt remains unaffected [60]. This increase in stem cell division rate translates to a 60% increase in TA and FD cell populations in our model, and thus a ∼ 60% increase in total crypt size. This prediction is close to the ∼ 65% increase in crypt area observed experimentally [22] for human hyperplastic polyps, and the ∼ 80% increase in the number of proliferating cells per crypt reported [60] in *Kras*-mutant crypts.

*APC* inactivation has been shown to result in the overpopulation of stem cells in the crypt [19, 27, 62, 63], without an increase in the stem cell division rate [62, 64]. By employing measurements from experiments and a model for stem cell dynamics, it has been inferred that crypts with *APC* inactivation have, on average, 1.6 times more stem cells compared to wild type crypts [27, 62]. Thus, for crypts with *APC* inactivation, we consider ∼ 28 stem cells. In our model, the average numbers of TA, FD and total cells per crypt are proportional to the number of stem cells, and thus a 60% increase in stem cell numbers results in 60% increase in the size of TA and FD cell compartments, as well as the total number of cells in a crypt. Experimentally, a proportional increase up to 100% has been observed in both the TA compartment size and total crypt size as a result of *Apc* inactivation [65].

Last, it has been experimentally observed [22] that the difference in the average number of mitoses per area between healthy human crypts and crypts from hyperplastic polyps and adenomas is not significant. Non-elevated mitotic index has also been reported for *Apc*-deficient crypts [65]. The average number of mitoses per area is expressed equivalently as the average ratio of proliferating cells to total number of cells, which, in our model, is approximated by *m*_TA_*/N*_tot_ ≈ *m*_TA_*/*(*m*_TA_ + *m*_FD_) = *γ/*(*d* + *γ*). Since the values for *d* and *γ* remain unchanged, our model predicts the same number of mitoses per area for both healthy and mutant crypts, in agreement with the experimental results.

## 5 Discussion

In this work, we present a multitype branching process model of cell population dynamics in colonic crypts. The simplicity of model formulation, along with its analytically obtained exact solution, provides insight into the the process by which both healthy and mutated colonic crypts maintain their size. For our model calibration, the only parameter fitting performed is for TA cell differentiation rate *d*; the remaining parameter values are estimated directly from the relevant literature. More specifically, we estimate *d* so that the average crypt size predicted by our model coincides with the experimental data. It is noteworthy that this fitting based only on average crypt size results in the predicted probability distributions for TA, FD and total cell populations to agree with experimental data.

Furthermore, the model’s stationarity condition for finite crypt size is simple, *d > λ*, and can also be interpreted as a criterion for malignancy in the process of colorectal carcinogenesis; our model predicts unbounded growth in cell numbers when the acquired mutations result in unregulated divisions of TA cells, with division rate *λ* exceeding their differentiation rate *d*. Indeed, in this model, homeostasis is achieved by adjusting the differentiation rate of TA cells. Another aspect of this homeostatic mechanism is that, for varying apoptosis rates, the crypt size distribution stays the same by adjusting the value of *d*.

Condition *d > λ* is also satisfied by the crypts in premalignant adenomas: In adenomatous crypts, the disruption of homeostasis results in larger yet finite crypt sizes. However, since adenomas undergo slow clonal expansions, possibly via crypt fission [1], adenomatous crypts can be best described not as stationary but as quasi-static. We should note that, in its present form, this model does not account for the crypt fission and loss; the incorporation of crypt fission and loss is an important direction for future work towards a better description of the expansion of the premalignant adenomas.

A limitation of our model is the assumption that the number *N*_0_ of stem cells is the same in all crypts. We note that values of 15 to 20 stem cells per crypt have been reported [44, 37]. Similarly, in murine crypts, a variation of ∼ 14% in stem cell number from crypt to crypt has been reported [38]. We compare our model results with constant *N*_0_ = 18 for all crypts, to results where *N*_0_ is allowed to randomly vary from crypt to crypt, with *N*_0_ = 15 *−* 21. The averaged cell population distributions for varying *N*_0_ coincide with the respective distributions for constant *N*_0_ (Supplemental Material, Fig. S4).

Another limitation of the model is that it presupposes the existence of homeostatic mechanisms responsible for maintaining the number of stem cells in the crypt [38, 25, 39]; thus, it is not an adequate model for crypt cell dynamics under changes in stem cell population, such as the replenishment of their numbers after an acute stem cell loss. It has been observed experimentally [8, 66] that stem cell homeostasis is based on feedback mechanisms, such as the plasticity of early TA cells to obtain stemness again if needed. However, a crypt cell population model that incorporates these mechanisms would be more complex, and it is outside of the scope of the present work.

## Data Accessibility

The code and data used to generate the figures of the main paper and Supplemental Material can be downloaded from GitHub at https://github.com/kmamis/stochastic_model_colonic_crypts. In this repository, there is also MATLAB Application “distribution calculator.mlapp” that takes as imput the stem cell number, division rates for stem and TA cells, FD cells apoptosis rate and the average number of cells per crypt, plots the cell population distributions predicted by the model, and calculates the average number of TA, FD cells per crypt, CV% of TA, FD and total cell populations, as well as the correlation coefficient between TA and FD cells.

## Authors’ Contributions

K.M. and I.B. designed research; K.M., R.Z. and I.B. performed research; K.M. and I.B. wrote the paper.

## Acknowledgements

We thank Simon Zhang, Applied Math student at the University of Washington, for proofreading the mathematical derivations of the Supplemental Material.

## References

[1] Zeki SS, Graham TA, Wright NA. 2011 Stem cells and their implications for colorectal cancer. Nature Reviews Gastroenterology and Hepatology 8, 90–100. (10.1038/nrgastro.2010.211)

[2] Yang J, Axelrod DE, Komarova NL. 2017 Determining the control networks regulating stem cell lineages in colonic crypts. Journal of Theoretical Biology 429, 190–203. (10.1016/j.jtbi.2017.06.033)

[3] Bravo R, Axelrod DE. 2013 A calibrated agent-based computer model of stochastic cell dynamics in normal human colon crypts useful for in silico experiments. Theoretical Biology and Medical Modelling 10, 66. (10.1186/1742-4682-10-66)

[4] Johnston MD, Edwards CM, Bodmer WF, Maini PK, Chapman SJ. 2007 Mathematical modeling of cell population dynamics in the colonic crypt and in colorectal cancer. Proceedings of the National Academy of Sciences 104, 4008–4013. (10.1073/pnas.0611179104)

[5] Hall PA, Coates PJ, Ansari B, Hopwood D. 1994 Regulation of cell number in the mammalian gastrointestinal tract: the importance of apoptosis. Journal of Cell Science 107, 3569–3577. (10.1242/jcs.107.12.3569)

[6] Barker N, Ridgway RA, Van Es JH, Van De Wetering M, Begthel H, Van Den Born M, Danenberg E, Clarke AR, Sansom OJ, Clevers H. 2009 Crypt stem cells as the cells-of-origin of intestinal cancer. Nature 457, 608–611. (10.1038/nature07602)

[7] Cancedda R, Mastrogiacomo M. 2023 Transit Amplifying Cells (TACs): a still not fully understood cell population. (10.3389/fbioe.2023.1189225)

[8] Clevers H. 2013 The intestinal crypt, a prototype stem cell compartment. Cell 154, 274–284. (10.1016/j.cell.2013.07.004)

[9] Nicolas P, Kim KM, Shibata D, Tavaré S. 2007 The stem cell population of the human colon crypt: Analysis via methylation patterns. PLoS Computational Biology 3, 0364–0374. (10.1371/journal.pcbi.0030028)

[10] Spit M, Koo BK, Maurice MM. 2018 Tales from the crypt: Intestinal niche signals in tissue renewal, plasticity and cancer. Open Biology 8, 180120. (10.1098/rsob.180120)

[11] Sanman LE, Chen IW, Bieber JM, Steri V, Trentesaux C, Hann B, Klein OD, Wu LF, Altschuler SJ. 2021 Transit-Amplifying cells coordinate changes in intestinal epithelial cell-type composition. Developmental Cell 56, 356–365. (10.1016/j.devcel.2020.12.020)

[12] Giles RH, Van Es JH, Clevers H. 2003 Caught up in a Wnt storm: Wnt signaling in cancer. Biochimica et Biophysica Acta - Reviews on Cancer 1653, 1–24. (10.1016/S0304-419X(03)00005-2)

[13] Van Leeuwen IMM, Byrne HM, Jensen OE, King JR. 2006 Crypt dynamics and colorectal cancer: advances in mathematical modelling. Cell Prolif 39, 157–181.

[14] Carulli AJ, Samuelson LC, Schnell S. 2014 Unraveling intestinal stem cell behavior with models of crypt dynamics. Integrative Biology 6, 243–257. (10.1039/c3ib40163d)

[15] Totafurno J, Bjerknes M, Cheng H. 1988 Variation in crypt size and its influence on the analysis of epithelial cell proliferation in the intestinal crypt. Biophysical Journal 54, 845–858. (10.1016/S0006-3495(88)83021-2)

[16] Gerike TG, Paulus U, Potten CS, Loeffler M. 1998 A dynamic model of proliferation and differentiation in the intestinal crypt based on a hypothetical intraepithelial growth factor. Cell Proliferation 31, 93–110. (10.1046/j.1365-2184.1998.00113.x)

[17] Meineke FA, Potten CS, Loeffler M. 2001 Cell migration and organization in the intestinal crypt using a lattice-free model. Cell Proliferation 34, 253–266. (10.1046/j.0960-7722.2001.00216.x)

[18] Van Leeuwen IM, Mirams GR, Walter A, Fletcher A, Murray P, Osborne J, Varma S, Young SJ, Cooper J, Doyle B, Pitt-Francis J, Momtahan L, Pathmanathan P, Whiteley JP, Chapman SJ, Gavaghan DJ, Jensen OE, King JR, Maini PK, Waters SL, Byrne HM. 2009 An integrative computational model for intestinal tissue renewal. Cell Proliferation 42, 617–636. (10.1111/j.1365-2184.2009.00627.x)

[19] Boman BM, Fields JZ, Bonham-Carter O, Runquist OA. 2001 Computer modeling implicates stem cell overproduction in colon cancer initiation. Cancer Research 61, 8408–8411.

[20] Dingli D, Traulsen A, Pacheco JM. 2007 Compartmental architecture and dynamics of hematopoiesis. PLoS ONE 2, e345. (10.1371/journal.pone.0000345)

[21] Lo WC, Chou CS, Gokoffski KK, Wan FY, Lander AD, Calof AL, Nie Q. 2009 Feedback regulation in multistage cell lineages. Mathematical Biosciences and Engineering 6, 59–82. (10.3934/mbe.2009.6.59)

[22] Wong WM, Mandir N, Goodlad RA, Wong BC, Garcia SB, Lam SK, Wright NA. 2002 Histogenesis of human colorectal adenomas and hyperplastic polyps: The role of cell proliferation and crypt fission. Gut 50, 212–217. (10.1136/gut.50.2.212)

[23] Tomlinson IPM, Bodmer WF. 1995 Failure of programmed cell death and differentiation as causes of tumors: Some simple mathematical models (tumorigenesis/apoptosis). Proceedings of the National Academy of Sciences 92, 11130–11134. (10.1073/pnas.92.24.11130)

[24] Paulus U, Potten CS, Loeffler M. 1992 A model of the control of cellular regeneration in the intestinal crypt after perturbation based solely on local stem cell regulation. Cell Prolif 25, 559–578. (10.1111/j.1365-2184.1992.tb01460.x)

[25] Snippert HJ, Schepers AG, Van Es JH, Simons BD, Clevers H. 2014 Biased competition between Lgr5 intestinal stem cells driven by oncogenic mutation induces clonal expansion. EMBO Reports 15, 62–69. (10.1002/embr.201337799)

[26] Bozic I. 2022 Quantification of the selective advantage of driver mutations is dependent on the underlying model and stage of tumor evolution. Cancer Research 82, 21–24. (10.1158/0008-5472.CAN-21-1064)

[27] Baker AM, Cereser B, Melton S, Fletcher AG, Rodriguez-Justo M, Tadrous PJ, Humphries A, Elia G, McDonald SA, Wright NA, Simons BD, Jansen M, Graham TA. 2014 Quantification of crypt and stem cell evolution in the normal and neoplastic human colon. Cell Reports 8, 940–947. (10.1016/j.celrep.2014.07.019)

[28] Kimmel M, Axelrod DE. 2002 Branching Processes in Biology. New York Berlin Heidelberg: Springer.

[29] Zhang R, Ukogu OA, Bozic I. 2023 Waiting times in a branching process model of colorectal cancer initiation. Theoretical Population Biology 151, 44–63. (10.1016/j.tpb.2023.04.001)

[30] Bozic I, Wu CJ. 2020 Delineating the evolutionary dynamics of cancer from theory to reality. Nature cancer 1, 580–588. (10.1038/s43018-020-0079-6)

[31] Bozic I, Antal T, Ohtsuki H, Carter H, Kim D, Chen S, Karchin R, Kinzler KW, Vogelstein B, Nowak MA. 2010 Accumulation of driver and passenger mutations during tumor progression. Proceedings of the National Academy of Sciences 107, 18545–18550. (10.1073/pnas.1010978107)

[32] Antal T, Krapivsky PL. 2011 Exact solution of a two-type branching process: Models of tumor progression. Journal of Statistical Mechanics: Theory and Experiment 2011, P08018. (10.1088/1742-5468/2011/08/P08018)

[33] Avanzini S, Kurtz DM, Chabon JJ, Moding EJ, Hori SS, Gambhir SS, Alizadeh AA, Diehn M, Reiter JG. 2020 A mathematical model of ctDNA shedding predicts tumor detection size. Sci. Adv 6, eabc4308. (10.1126/sciadv.abc4308)

[34] Clayton E, Doupé DP, Klein AM, Winton DJ, Simons BD, Jones PH. 2007 A single type of progenitor cell maintains normal epidermis. Nature 446, 185–189. (10.1038/nature05574)

[35] Antal T, Krapivsky PL. 2010 Exact solution of a two-type branching process: Clone size distribution in cell division kinetics. Journal of Statistical Mechanics: Theory and Experiment 2010, P07028. (10.1088/1742-5468/2010/07/P07028)

[36] Shahriyari L, Komarova NL. 2013 Symmetric vs. asymmetric stem cell divisions: an adaptation against cancer?. PloS one 8, e76195. (10.1371/journal.pone.0076195)

[37] Gehart H, Clevers H. 2019 Tales from the crypt: New insights into intestinal stem cells. Nature Reviews Gastroenterology and Hepatology 16, 19–34. (10.1038/s41575-018-0081-y)

[38] Snippert HJ, van der Flier LG, Sato T, van Es JH, van den Born M, Kroon-Veenboer C, Barker N, Klein AM, van Rheenen J, Simons BD, Clevers H. 2010 Intestinal crypt homeostasis results from neutral competition between symmetrically dividing Lgr5 stem cells. Cell 143, 134–144. (10.1016/j.cell.2010.09.016)

[39] Lopez-Garcia C, Klein AM, Simons BD, Winton DJ. 2010 Intestinal stem cell replacement follows a pattern of neutral drift. Science 330, 822–825. (10.1126/science.1196236)

[40] Leblond CP. 1964 Classification of cell populations on the basis of their proliferative behavior. In Congdon CC, Mori-Chavez P, editors, Control of Cell Division and the Induction of Cancer, pp. 119–150. Bethesda, MD: National Cancer Institute.

[41] Dewanji A, Luebeck EG, Moolgavkar SH. 2005 A generalized Luria-Delbrück model. Mathematical Biosciences 197, 140–152. (10.1016/j.mbs.2005.07.003)

[42] Darwich AS, Aslam U, Ashcroft DM, Rostami-Hodjegan A. 2014 Meta-analysis of the turnover of intestinal epithelia in preclinical animal species and humans. (10.1124/dmd.114.058404)

[43] Potten CS, Loeffler M. 1990 Stem cells: attributes, cycles, spirals, pitfalls and uncertainties Lessons for and from the Crypt. Development 110, 1001–1020. (10.1242/dev.110.4.1001)

[44] Gasnier M, Lim HYG, Barker N. 2023 Role of Wnt signaling in the maintenance and regeneration of the intestinal epithelium. Current Topics in Developmental Biology 153, 281–326. (10.1016/bs.ctdb.2023.01.001)

[45] Marshman E, Booth C, Potten CS. 2002 The intestinal epithelial stem cell. BioEssays 24, 91–98. (10.1002/bies.10028)

[46] Shahriyari L, Komarova NL, Jilkine A. 2016 The role of cell location and spatial gradients in the evolutionary dynamics of colon and intestinal crypts. Biology Direct 11, 42. (10.1186/s13062-016-0141-6)

[47] Potten CS, Kellett M, Roberts SA, Rew DA, Wilson GD. 1992a Measurement of in vivo proliferation in human colorectal mucosa using bromodeoxyuridine. Gut 33, 71–78. (10.1136/gut.33.1.71)

[48] Potten CS, Kellett M, Rew DA, Roberts SA. 1992b Proliferation in human gastrointesinal epithelium using bromodeoxyuridine in vivo: Data for different sites, proximity to a tumour, and polyposis coli. Gut 33, 524–529. (10.1136/gut.33.4.524)

[49] Roncucci L, de Leon MP, Scalmati A, Malagoli G, Pratissoli S, Perini M, Chahin NJ. 1988 The influence of age on colonic epithelial cell proliferation. Cancer 62, 2373–2377. (10.1002/1097-0142(19881201)62:11¡2373::aid-cncr2820621120¿3.0.co;2-y)

[50] Tomlinson I, Sasieni P, Bodmer W. 2002 How many mutations in a cancer?. American Journal of Pathology 160, 755–758. (10.1016/S0002-9440(10)64896-1)

[51] Bjerknes M, Cheng H. 1981 The stem-cell zone of the small intestinal epithelium. IV. Effects of resecting 30% of the small intestine. The American Journal of Anatomy 160, 93–103. (10.1002/aja.1001600108)

[52] Miller I, Min M, Yang C, Tian C, Gookin S, Carter D, Spencer SL. 2018 Ki67 is a graded rather than a binary marker of proliferation versus quiescence. Cell Reports 24, 1105–1112. (10.1016/j.celrep.2018.06.110)

[53] Gabbutt C, Schenck RO, Weisenberger DJ, Kimberley C, Berner A, Househam J, Lakatos E, Robertson-Tessi M, Martin I, Patel R, Clark SK, Latchford A, Barnes CP, Leedham SJ, Anderson AR, Graham TA, Shibata D. 2022 Fluctuating methylation clocks for cell lineage tracing at high temporal resolution in human tissues. Nature Biotechnology 40, 720–730. (10.1038/s41587-021-01109-w)

[54] Nicholson AM, Olpe C, Hoyle A, Thorsen AS, Rus T, Colombé M, Brunton-Sim R, Kemp R, Marks K, Quirke P, Malhotra S, ten Hoopen R, Ibrahim A, Lindskog C, Myers MB, Parsons B, Tavaré S, Wilkinson M, Morrissey E, Winton DJ. 2018 Fixation and Spread of Somatic Mutations in Adult Human Colonic Epithelium. Cell Stem Cell 22, 909–918. (10.1016/j.stem.2018.04.020)

[55] Lamlum H, Papadopoulou A, Ilyas M, Rowan A, Gillet C, Hanby A, Talbot I, Bodmer W Tomlinson 2000 APC mutations are sufficient for the growth of early colorectal adenomas. Proceedings of the National Academy of Sciences 97, 2225–2228. (10.1073/pnas.040564697)

[56] van Neerven SM, de Groot NE, Nijman LE, Scicluna BP, van Driel MS, Lecca MC, Warmerdam DO, Kakkar V, Moreno LF, Vieira Braga FA, Sanches DR, Ramesh P, ten Hoorn S, Aelvoet AS, van Boxel MF, Koens L, Krawczyk PM, Koster J, Dekker E, Medema JP, Winton DJ, Bijlsma MF, Morrissey E, Léveillé N, Vermeulen L. 2021 Apc-mutant cells act as supercompetitors in intestinal tumour initiation. Nature 594, 436–441. (10.1038/s41586-021-03558-4)

[57] Dow LE, O’Rourke KP, Simon J, Tschaharganeh DF, Van Es JH, Clevers H, Lowe SW. 2015 Apc restoration promotes cellular differentiation and reestablishes crypt homeostasis in colorectal cancer. Cell 161, 1539–1552. (10.1016/j.cell.2015.05.033)

[58] Boman BM, Walters R, Fields JZ, Kovatich AJ, Zhang T, Isenberg GA, Goldstein SD, Palazzo JP. 2004 Colonic crypt changes during adenoma development in familial adenomatous polyposis. Immuno-histochemical evidence for expansion of the crypt base cell population. Am J Pathol 165, 1489–1498. (10.1016/S0002-9440(10)63407-4)

[59] Paterson C, Clevers H, Bozic I. 2020 Mathematical model of colorectal cancer initiation. Proceedings of the National Academy of Sciences 117, 20681–20688. (10.1073/pnas.2003771117)

[60] Feng Y, Bommer GT, Zhao J, Green M, Sands E, Zhai Y, Brown K, Burberry A, Cho KR, Fearon ER. 2011 Mutant kras promotes hyperplasia and alters differentiation in the colon epithelium but does not expand the presumptive stem cell pool. Gastroenterology 141, 1003–1013. (10.1053/j.gastro.2011.05.007)

[61] Chan TL, Zhao W, Leung SY, Yuen ST. 2003 BRAF and KRAS mutations in colorectal hyperplastic polyps and serrated adenomas. Cancer Research 63, 4878–4881.

[62] Kozar S, Morrissey E, Nicholson AM, van der Heijden M, Zecchini HI, Kemp R, Tavaré S, Vermeulen L, Winton DJ. 2013 Continuous clonal labeling reveals small numbers of functional stem cells in intestinal crypts and adenomas. Cell stem cell 13, 626–633. (10.1016/j.stem.2013.08.001)

[63] Zhang T, Ahn K, Emerick B, Modarai SR, Opdenaker LM, Palazzo J, Schleiniger G, Fields JZ, Boman BM. 2020 APC mutations in human colon lead to decreased neuroendocrine maturation of ALDH+ stem cells that alters GLP-2 and SST feedback signaling: Clue to a link between WNT and retinoic acid signalling in colon cancer development. PLoS ONE 15, e0239601. (10.1371/journal.pone.0239601)

[64] Bruschi M, Garnier L, Cleroux E, Giordano A, Dumas M, Bardet AF, Kergrohen T, Quesada S, Cesses P, Weber M, Gerbe F, Jay P. 2020 Loss of Apc rapidly impairs DNA methylation programs and cell fate decisions in Lgr5+ intestinal stem cells. Cancer Research 80, 2101–2113. (10.1158/0008-5472.CAN-19-2104)

[65] Sansom OJ, Meniel VS, Muncan V, Phesse TJ, Wilkins JA, Reed KR, Vass JK, Athineos D, Clevers H, Clarke AR. 2007 Myc deletion rescues Apc deficiency in the small intestine. Nature 446, 676–679. (10.1038/nature05674)

[66] Clevers H, Loh KM, Nusse R. 2014 An integral program for tissue renewal and regeneration: Wnt signaling and stem cell control. Science 346. (10.1126/science.1248012)

