## Supplemental Material for "Stochastic model for cell population dynamics quantifies homeostasis in colonic crypts and its disruption in early tumorigenesis"

#### S1 Branching process and its forward Kolmogorov equation

In the present work, we propose the following continuous-time, constant-rate, three-type branching process for the population dynamics of stem (S), transit-amplifying (TA), and fully differentiated (FD) cells in colonic crypts:

$$\begin{aligned} &\text{S} \xrightarrow{r} \text{S TA} \\ &\text{TA} \xrightarrow{\lambda} \text{TA TA} \\ &\text{TA} \xrightarrow{d} \text{FD} \\ &\text{FD} \xrightarrow{\gamma} \emptyset. \end{aligned} \tag{S1}$$

In Eq. (S1), stem cells divide asymmetrically with rate $r$, TA cells divide symmetrically with rate $\lambda$ or differentiate into FD cells with rate $d$, and FD cells undergo apoptosis with rate $\gamma$.

Having the rate equations (S1) of the branching process, we formulate the forward Kolmogorov equation [1, Sec. 5.6] for the probability $P_{n_0,n_1,n_2}(t)$ of having $n_0$ stem cells, $n_1$ TA cells, and $n_2$ FD cells at time $t$:

$$\begin{aligned} \frac{\mathrm{d}P_{n_0,n_1,n_2}(t)}{\mathrm{d}t} &= rN_0[P_{n_0,n_1-1,n_2}(t) - P_{n_0,n_1,n_2}(t)] + \lambda(n_1-1)P_{n_0,n_1-1,n_2}(t) + \\ &+ d(n_1+1)P_{n_0,n_1+1,n_2-1} - (\lambda+d)n_1 P_{n_0,n_1,n_2}(t) + \gamma(n_2+1)P_{n_0,n_1,n_2+1}(t) - \gamma n_2 P_{n_0,n_1,n_2}(t). \end{aligned} \tag{S2}$$

Eq. (S2) describes the stochastic expansion of cell populations in a colonic crypt from stem cells to its steady-steady size. Thus, as its initial condition, we choose

$$P_{n_0,n_1,n_2}(0) = \delta_{n_0,N_0}\delta_{n_1,0}\delta_{n_2,0}, \tag{S3}$$

where $\delta_{n,k}$ is Kronecker's delta, modeling the fact that at initial time, crypts contain only $N_0$ stem cells. We also note, that, since stem cells only divide asymmetrically, their initial population is maintained and does not fluctuate stochastically; in every time instance, there are $N_0$ stem cells per crypt. This results in the simplification of the joint probability of cell populations $P_{n_0,n_1,n_2}(t) = \delta_{n_0,N_0}P_{n_1,n_2}(t)$. Thus Eqs. (S2),(S3), are expressed for the joint probability of TA and FD cell populations $P_{n_1,n_2}(t)$ as

$$\begin{aligned} \frac{\mathrm{d}P_{n_1,n_2}(t)}{\mathrm{d}t} &= rN_0[P_{n_1-1,n_2}(t) - P_{n_1,n_2}(t)] + \lambda(n_1-1)P_{n_1-1,n_2}(t) + d(n_1+1)P_{n_1+1,n_2-1}(t) - \\ &- (\lambda+d)n_1 P_{n_1,n_2}(t) + \gamma(n_2+1)P_{n_1,n_2+1}(t) - \gamma n_2 P_{n_1,n_2}(t), \quad P_{n_1,n_2}(0) = \delta_{n_1,0}\delta_{n_2,0}. \end{aligned} \tag{S4}$$

#### S2 Derivation of ODEs for average sizes of cell populations

We introduce the average population sizes

$$m_{\mathrm{TA}}(t) = \sum_{n_1,n_2\geq 0} n_1 P_{n_1,n_2}(t), \quad m_{\mathrm{FD}}(t) = \sum_{n_1,n_2\geq 0} n_2 P_{n_1,n_2}(t),$$

and the second order moments

$$R_{\mathrm{TA}}(t) = \sum_{n_1,n_2\geq 0} n_1^2 P_{n_1,n_2}(t), \quad R_{\mathrm{FD}}(t) = \sum_{n_1,n_2\geq 0} n_2^2 P_{n_1,n_2}(t), \quad R_{\mathrm{TA,FD}}(t) = \sum_{n_1,n_2\geq 0} n_1 n_2 P_{n_1,n_2}(t).$$

For the derivation of the ODE for $m_{\mathrm{TA}}(t)$, we multiply both sides of Eq. (S4) by $n_1$ and summing over all $n_1, n_2 \geq 0$, under the convention that $P_{n_1,n_2}(t) \equiv 0$ for $n_1 < 0$ or $n_2 < 0$. For each term of Eq. (S4), we calculate:

- $rN_0 \sum_{n_1,n_2} n_1 P_{n_1-1,n_2}(t) = rN_0 \left( \sum_{n_1,n_2} (n_1-1) P_{n_1-1,n_2}(t) + \sum_{n_1,n_2} P_{n_1-1,n_2}(t) \right) = rN_0\,[m_{\mathrm{TA}}(t)+1],$
- $rN_0 \sum_{n_1,n_2} n_1 P_{n_1,n_2}(t) = rN_0 m_{\mathrm{TA}}(t),$
- $\lambda \sum_{n_1,n_2} n_1(n_1-1) P_{n_1-1,n_2}(t) = \lambda \left( \sum_{n_1,n_2} (n_1-1)^2 P_{n_1-1,n_2}(t) + \sum_{n_1,n_2} (n_1-1) P_{n_1-1,n_2}(t) \right) =$
$= \lambda\,[R_{\mathrm{TA}}(t) + m_{\mathrm{TA}}(t)],$
- $d \sum_{n_1,n_2} n_1(n_1+1) P_{n_1+1,n_2-1}(t) = d \left( \sum_{n_1,n_2} (n_1+1)^2 P_{n_1+1,n_2-1}(t) - \sum_{n_1,n_2} (n_1+1) P_{n_1+1,n_2-1}(t) \right) =$
$= d\,[R_{\mathrm{TA}}(t) - m_{\mathrm{TA}}(t)],$
- $(\lambda+d) \sum_{n_1,n_2} n_1^2 P_{n_1,n_2}(t) = (\lambda+d) R_{\mathrm{TA}}(t),$
- $\gamma \sum_{n_1,n_2} n_1(n_2+1) P_{n_1,n_2+1}(t) = \gamma R_{\mathrm{TA,FD}}(t),$
- $\gamma \sum_{n_1,n_2} n_1 n_2 P_{n_1,n_2}(t) = \gamma R_{\mathrm{TA,FD}}(t).$

Under the above calculations, Eq. (S4) results in

$$\frac{\mathrm{d}m_{\mathrm{TA}}(t)}{\mathrm{d}t} = rN_0\,[\cancel{m_{\mathrm{TA}}(t)}+1] - \cancel{rN_0 m_{\mathrm{TA}}(t)} + \lambda\,[\cancel{R_{\mathrm{TA}}(t)} + m_{\mathrm{TA}}(t)] + d\,[\cancel{R_{\mathrm{TA}}(t)} - m_{\mathrm{TA}}(t)] - \\ - \cancel{(\lambda+d)R_{\mathrm{TA}}(t)} + \cancel{\gamma R_{\mathrm{TA,FD}}(t)} - \cancel{\gamma R_{\mathrm{TA,FD}}(t)}, \quad m_{\mathrm{TA}}(0) = 0. \tag{S5}$$

Similarly, for the derivation of the ODE for $m_{\mathrm{FD}}(t)$, we multiply both sides of Eq. (S4) by $n_2$ and summing over all $n_1, n_2 \geq 0$, under the convention that $P_{n_1,n_2}(t) \equiv 0$ for $n_1 < 0$ or $n_2 < 0$. For each term of Eq. (S4), we calculate:

- $rN_0 \left( \sum_{n_1,n_2} n_2 P_{n_1-1,n_2}(t) - \sum_{n_1,n_2} n_2 P_{n_1,n_2}(t) \right) = rN_0\,[m_{\mathrm{FD}}(t) - m_{\mathrm{FD}}(t)] = 0,$
- $\lambda \sum_{n_1,n_2} (n_1-1) n_2 P_{n_1-1,n_2}(t) = \lambda R_{\mathrm{TA,FD}}(t),$
- $d \sum_{n_1,n_2} (n_1+1) n_2 P_{n_1+1,n_2-1}(t) = d \left( \sum_{n_1,n_2} (n_1+1)(n_2-1) P_{n_1+1,n_2-1}(t) + \sum_{n_1,n_2} (n_1+1) P_{n_1+1,n_2-1}(t) \right) =$
$= d\,[R_{\mathrm{TA,FD}}(t) + m_{\mathrm{TA}}(t)],$
- $(\lambda+d) \sum_{n_1,n_2} n_1 n_2 P_{n_1,n_2}(t) = (\lambda+d) R_{\mathrm{TA,FD}}(t),$
- $\gamma \sum_{n_1,n_2} n_2(n_2+1) P_{n_1,n_2+1}(t) = \gamma \left( \sum_{n_1,n_2} (n_2+1)^2 P_{n_1,n_2+1}(t) - \sum_{n_1,n_2} (n_2+1) P_{n_1,n_2+1}(t) \right) =$
$= \gamma\,[R_{\mathrm{FD}}(t) - m_{\mathrm{FD}}(t)],$

- $\gamma \sum_{n_1,n_2} n_2^2 P_{n_1,n_2}(t) = \gamma R_{\mathrm{FD}}(t).$

Under the above calculations, Eq. (S4) results in

$$\begin{aligned}\frac{\mathrm{d}m_{\mathrm{FD}}(t)}{\mathrm{d}t} &= \cancel{\lambda R_{\mathrm{TA,FD}}(t)} + d\left[\cancel{R_{\mathrm{TA,FD}}(t)} + m_{\mathrm{TA}}(t)\right] - \\ &- \cancel{(\lambda+d)R_{\mathrm{TA,FD}}(t)} + \gamma\left[\cancel{R_{\mathrm{FD}}(t)} - m_{\mathrm{FD}}(t)\right] - \cancel{\gamma R_{\mathrm{FD}}(t)}, \quad m_{\mathrm{FD}}(0) = 0. \end{aligned} \tag{S6}$$

Eqs. (S5), (S6) are the system of equations (2.2) of the main paper.

### S3 Formulation and solution of the PGF equation for TA cell population

The branching process for the TA cell population only reads:

$$\begin{aligned} &\emptyset \xrightarrow{rN_0} \mathrm{TA} \\ &\mathrm{TA} \xrightarrow{\lambda} \mathrm{TA}\ \mathrm{TA} \\ &\mathrm{TA} \xrightarrow{d} \emptyset. \end{aligned} \tag{S7}$$

The forward Kolmogorov equation that corresponds to branching process (S7) is

$$\frac{\mathrm{d}P_{n_1}(t)}{\mathrm{d}t} = rN_0 P_{n_1-1}(t) + \lambda(n_1-1)P_{n_1-1}(t) + d(n_1+1)P_{n_1+1}(t) - [rN_0 + (\lambda+d)n_1]P_{n_1}(t), \quad P_{n_1}(0) = \delta_{n_1,0}. \tag{S8}$$

Initial condition of Eq. (S8) describes that there are no TA cells at $t = 0$. Eq. (S8) is recast in a more amenable form by considering the probability-generating function (PGF) $F_{\mathrm{TA}}(x,t) = \sum_{n_1 \geq 0} x^{n_1} P_{n_1}(t)$, with $|x| \leq 1$. By multiplying both sides of Eq. (S8) with $x^{n_1}$, and summing over all $n_1 \geq 0$, we obtain

$$\partial_t F_{\mathrm{TA}}(x,t) = rN_0(x-1)F_{\mathrm{TA}}(x,t) + [\lambda x^2 + d - (\lambda+d)x]\partial_x F_{\mathrm{TA}}(x,t), \quad F_{\mathrm{TA}}(x,0) = 1, \tag{S9}$$

where $\partial_t$, $\partial_x$ denote the partial derivatives with respect to $t$, $x$. The characteristic equation, see e.g. [2, Sec. 8.7], [3, Sec. 2.1], of Eq. (S9) is

$$\frac{\mathrm{d}t}{1} = \frac{-\mathrm{d}x}{\lambda x^2 + d - (\lambda+d)x} = \frac{\mathrm{d}F_{\mathrm{TA}}}{rN_0(x-1)F_{\mathrm{TA}}}.$$

The first ODE to solve is

$$\frac{\mathrm{d}t}{1} = \frac{-\mathrm{d}x}{\lambda x^2 + d - (\lambda+d)x}$$

whose solution reads

$$\frac{e^{(d-\lambda)t}(d-\lambda x)}{1-x} = C. \tag{S10}$$

The second ODE is

$$\frac{\mathrm{d}F_{\mathrm{TA}}}{F_{\mathrm{TA}}} = -rN_0 \frac{x-1}{\lambda x^2 + d - (\lambda+d)x}\mathrm{d}x,$$

whose solution, under the condition $d > \lambda$, reads

$$F_{\mathrm{TA}}(d-\lambda x)^{rN_0/\lambda} = C_1. \tag{S11}$$

Thus, for Eq. (S9), we obtain the general solution

$$F_{\mathrm{TA}}(d-\lambda x)^{rN_0/\lambda} = \Psi\left(\frac{e^{(d-\lambda)t}(d-\lambda x)}{1-x}\right). \tag{S12}$$

Function $\Psi$ is determined by the initial condition $F_{\mathrm{TA}}(t=0) = 1$, resulting in

$$\Psi(C) = \left(d - \lambda\frac{d-C}{\lambda-C}\right)^{rN_0/\lambda}.$$

Thus, the solution to Eq. (S9) is determined to

$$F_{\mathrm{TA}}(x,t) = \left[ \left( d - \lambda \frac{d - \frac{e^{(d-\lambda)t}(d-\lambda x)}{1-x}}{\lambda - \frac{e^{(d-\lambda)t}(d-\lambda x)}{1-x}} \right) \Big/ (d-\lambda x) \right]^{rN_0/\lambda}. \tag{S13}$$

We calculate the steady-state PGF $F_{\mathrm{TA}}(x)$ by taking the limit $t \to \infty$ of Eq. (S13). For $d > \lambda$ the long-time limit results in the indeterminate form $\infty/\infty$, which we resolve by using de l'Hôpital's rule:

$$F_{\mathrm{TA}}(x) = \left( \frac{d-\lambda}{d-\lambda x} \right)^{rN_0/\lambda}. \tag{S14}$$

Stationary PGF, Eq. (S14), corresponds to a generalized negative binomial distribution, with the number of successes being $rN_0/\lambda$ and success probability $(d-\lambda)/d$. Thus, the steady-state probability mass function for $n_1 = 0, 1, 2, \ldots$ TA cells is given by the explicit formula

$$P_{n_1} = \frac{\Gamma(n_1 + (rN_0/\lambda))}{n_1! \Gamma(rN_0/\lambda)} \left( \frac{d-\lambda}{d} \right)^{rN_0/\lambda} \left( \frac{\lambda}{d} \right)^{n_1}, \tag{S15}$$

with $\Gamma(\cdot)$ being the gamma function. Eq. (S15) is Eq. (2.6) of the main paper.

### S4 Approximate steady-state distribution for FD cell population

Let us assume that the number $n_1$ of TA cells is constant. In this case, the branching process for the FD cell population only reads:

$$\begin{aligned} &\emptyset \xrightarrow{dn_1} \mathrm{FD} \\ &\mathrm{FD} \xrightarrow{\gamma} \emptyset. \end{aligned} \tag{S16}$$

That is, FD cells follow a death process with rate $\gamma$, under the immigration term $dn_1$. The forward Kolmogorov equation that corresponds to branching process (S16) is

$$\frac{\mathrm{d}P_{n_2}(t)}{\mathrm{d}t} = dn_1 P_{n_2-1} + \gamma(n_2+1)P_{n_2+1} - (dn_1 + \gamma n_2)P_{n_2}, \quad P_{n_2}(0) = \delta_{n_2,0}. \tag{S17}$$

As in the section S3, we recast Eq. (S17) into an equation for the PGF $F_{\mathrm{FD}}(x,t) = \sum_{n_2 \geq 0} x^{n_2} P_{n_2}(t)$:

$$\partial_t F_{\mathrm{FD}}(x,t) = dn_1(x-1)F_{\mathrm{FD}} - \gamma(x-1)\partial_x F_{\mathrm{FD}}, \quad F_{\mathrm{FD}}(x,0) = 1. \tag{S18}$$

The corresponding characteristic equation is

$$\frac{\mathrm{d}t}{1} = \frac{\mathrm{d}x}{\gamma(x-1)} = \frac{\mathrm{d}F_{\mathrm{FD}}}{dn_1(x-1)F_{\mathrm{FD}}}.$$

The first ODE to solve is

$$\frac{\mathrm{d}t}{1} = \frac{\mathrm{d}x}{\gamma(x-1)}$$

whose solution reads

$$\frac{e^{\gamma t}}{1-x} = C. \tag{S19}$$

The second ODE is

$$\frac{\mathrm{d}F_{\mathrm{FD}}}{F_{\mathrm{FD}}} = \frac{dn_1(x-1)}{\gamma(x-1)}\mathrm{d}x = \frac{dn_1}{\gamma}\mathrm{d}x$$

whose solution reads

$$F_{\mathrm{FD}} \exp\left( -\frac{dn_1}{\gamma} x \right) = C_1. \tag{S20}$$

Thus, for Eq. (S18), we obtain the general solution

$$F_{\mathrm{FD}} \exp\left( -\frac{dn_1}{\gamma} x \right) = \Psi\left( \frac{e^{\gamma t}}{1-x} \right). \tag{S21}$$

We determine function $\Psi$ by the initial condition $F_{\mathrm{FD}}(t=0)=1$, resulting in

$$\Psi(C) = \exp\left(-\frac{dn_1}{\gamma}\frac{C-1}{C}\right).$$

Thus, the solution to Eq. (S18) is determined to

$$F_{\mathrm{FD}}(x,t) = \exp\left(\frac{dn_1}{\gamma}\left[x - \left(\frac{e^{\gamma t}}{1-x} - 1\right)\Big/\frac{e^{\gamma t}}{1-x}\right]\right). \tag{S22}$$

We calculate the steady-state PGF $F_{\mathrm{FD}}(x,t)$ by taking the limit $t\to\infty$ of Eq. (S22), and using de l'Hôpital's rule:

$$F_{\mathrm{FD}}(x) = \exp\left(\frac{dn_1}{\gamma}(x-1)\right). \tag{S23}$$

PGF of Eq. (S23) correspond to a Poisson distribution with parameter $dn_1/\gamma$:

$$p(n_2;n_1) = \frac{(dn_1/\gamma)^{n_2}}{n_2!}e^{-dn_1/\gamma}, \quad n_2 = 0,1,2,\ldots \tag{S24}$$

Thus, for the case of constant number $n_1$ of TA cells, the number $n_2$ of FD cells follows the Poisson distribution, Eq. (S24). In our case where the number of TA cells is not constant, we approximate the probability $P_{n_2}$ of having $n_2$ FD cells in the steady state, by summing $p(n_2;n_1)$ for all $n_1$, each one weighted by the probability $P_{n_1}$, Eq. (S15), of having $n_1$ TA cells in the steady state:

$$P_{n_2} \approx \sum_{n_1=0}^{\infty} P_{n_1}p(n_2;n_1) = \frac{1}{\Gamma(rN_0/\lambda)}\left(\frac{d-\lambda}{d}\right)^{rN_0/\lambda}\sum_{n_1=0}^{\infty}\frac{\Gamma(n_1+(rN_0/\lambda))}{n_1!}\left(\frac{\lambda}{d}\right)^{n_1}\frac{(dn_1/\gamma)^{n_2}}{n_2!}e^{-dn_1/\gamma}. \tag{S25}$$

Eq. (S25) is Eq. (2.8) of the main paper. We easily implement Eq. (S25) using the functions `nbinpdf`, `poisspdf` in MATLAB, for negative binomial and Poisson distributions respectively.

### S5 Equation for the joint PGF of TA and FD cell populations

#### S5.1 Equation formulation

For the branching process (S1), we choose to formulate and solve the corresponding backward Kolmogorov equation. The same choice is made in Refs. [4, 5, 6], based on the observation that the backward equation is solved more easily than the forward one, see [2, Sec. 8.9]. For its formulation, we consider the probability $P^{(N_0,0,0)}_{n_0,n_1,n_2}(t)$ of having $n_0$ stem cells, $n_1$ TA cells and $n_2$ FD cells at time $t$ given that at initial time $t=0$ there are $N_0$ stem cells and no TA and FD cells. For $P^{(N_0,0,0)}_{n_0,n_1,n_2}(t)$, the backward Kolmogorov equation reads [1, Sec. 5.6]:

$$\frac{\mathrm{d}P^{(N_0,0,0)}_{n_0,n_1,n_2}(t)}{\mathrm{d}t} = rN_0P^{(N_0,1,0)}_{n_0,n_1,n_2}(t) - rN_0P^{(N_0,0,0)}_{n_0,n_1,n_2}(t), \quad P^{(N_0,0,0)}_{n_0,n_1,n_2}(0) = \delta_{n_0,N_0}\delta_{n_1,0}\delta_{n_2,0}. \tag{S26}$$

For the closure of Eq. (S26), we need the following auxiliary backward Kolmogorov equations with different initial conditions:

$$\frac{\mathrm{d}P^{(0,1,0)}_{n_0,n_1,n_2}(t)}{\mathrm{d}t} = \lambda P^{(0,2,0)}_{n_0,n_1,n_2}(t) + dP^{(0,0,1)}_{n_0,n_1,n_2}(t) - (\lambda+d)P^{(0,1,0)}_{n_0,n_1,n_2}(t), \quad P^{(0,1,0)}_{n_0,n_1,n_2}(0) = \delta_{n_0,0}\delta_{n_1,1}\delta_{n_2,0}, \tag{S27}$$

$$\frac{\mathrm{d}P^{(0,0,1)}_{n_0,n_1,n_2}(t)}{\mathrm{d}t} = \gamma\delta_{n_0,0}\delta_{n_1,0}\delta_{n_2,0} - \gamma P^{(0,0,1)}_{n_0,n_1,n_2}(t), \quad P^{(0,0,1)}_{n_0,n_1,n_2}(0) = \delta_{n_0,0}\delta_{n_1,0}\delta_{n_2,1}. \tag{S28}$$

We introduce the joint PGF for stem, TA and FD cell populations

$$G(x_0,x_1,x_2,t) = \sum_{n_0,n_1,n_2\geq 0} x_0^{n_0}x_1^{n_1}x_2^{n_2}P^{(N_0,0,0)}_{n_0,n_1,n_2}(t),$$

$$G_1(x_0,x_1,x_2,t) = \sum_{n_0,n_1,n_2\geq 0} x_0^{n_0}x_1^{n_1}x_2^{n_2}P^{(0,1,0)}_{n_0,n_1,n_2}(t),$$

$$G_2(x_0,x_1,x_2,t) = \sum_{n_0,n_1,n_2\geq 0} x_0^{n_0}x_1^{n_1}x_2^{n_2}P^{(0,0,1)}_{n_0,n_1,n_2}(t).$$

By multiplying both sides of each Eq. (S26)-(S28) by $x_0^{n_0}x_1^{n_1}x_2^{n_2}$ and summing over all $n_0, n_1, n_2 \geq 0$ we obtain equations for the PGFs. Also, we invoke the following independence arguments: Since the progenies of $N_0$ stem cells are independent from the progenies of 1 TA cell, the PGF corresponding to $P^{(N_0,1,0)}_{n_0,n_1,n_2}(t)$ is $G(x_0,x_1,x_2,t)\cdot G_1(x_0,x_1,x_2,t)$. Similarly, since the progenies of two TA cells are independent from each other, the PGF corresponding to $P^{(0,2,0)}_{n_0,n_1,n_2}(t)$ is $G_1^2(x_0,x_1,x_2,t)$. Thus, we obtain

$$\partial_t G = rN_0(G_1-1)G, \quad G(t=0) = x_0^{N_0}, \tag{S29}$$

$$\partial_t G_1 = \lambda G_1^2 + dG_2 - (\lambda+d)G_1, \quad G_1(t=0) = x_1, \tag{S30}$$

$$\partial_t G_2 = \gamma - \gamma G_2, \quad G_2(t=0) = x_2. \tag{S31}$$

We now use the fact that the number of stem cells stays constant and equal to their number at the initial time, in every realization of the branching process (S1); $P^{(N_0,0,0)}_{n_0,n_1,n_2}(t) = \delta_{n_0,N_0}P^{(0,0)}_{n_1,n_2}(t)$, $P^{(0,1,0)}_{n_0,n_1,n_2}(t) = \delta_{n_0,0}P^{(1,0)}_{n_1,n_2}(t)$, $P^{(0,0,1)}_{n_0,n_1,n_2}(t) = \delta_{n_0,0}P^{(0,1)}_{n_1,n_2}(t)$. Thus, by introducing the joint PGF for TA and FD cell populations only

$$F(x_1,x_2,t) = \sum_{n_1,n_2\geq 0} x_1^{n_1}x_2^{n_2}P^{(0,0)}_{n_1,n_2}(t),$$

$$F_1(x_1,x_2,t) = \sum_{n_1,n_2\geq 0} x_1^{n_1}x_2^{n_2}P^{(1,0)}_{n_1,n_2}(t),$$

$$F_2(x_1,x_2,t) = \sum_{n_1,n_2\geq 0} x_1^{n_1}x_2^{n_2}P^{(0,1)}_{n_1,n_2}(t),$$

the $G$-PGFs are expressed as

$$G(x_0,x_1,x_2,t) = x_0^{N_0}F(x_1,x_2,t), \quad G_1(x_0,x_1,x_2,t) = x_0^0F_1(x_1,x_2,t), \quad G_2(x_0,x_1,x_2,t) = x_0^0F_2(x_1,x_2,t).$$

This results in the simplification of Eqs. (S29)-(S31) into

$$\partial_t F = rN_0(F_1-1)F, \quad F(t=0) = 1, \tag{S32}$$

$$\partial_t F_1 = \lambda F_1^2 + dF_2 - (\lambda+d)F_1, \quad F_1(t=0) = x_1, \tag{S33}$$

$$\partial_t F_2 = \gamma - \gamma F_2, \quad F_2(t=0) = x_2. \tag{S34}$$

#### S5.2 Equation solution

Eq. (S34) is linear, and thus its solution reads:

$$F_2(t) = 1-(1-x_2)e^{-\gamma t}. \tag{S35}$$

Substituting solution (S35) into Eq. (S33), and by performing the change in temporal variable $s = e^{-\gamma t}$, we obtain the Riccati equation

$$\partial_s F_1(s) = -\frac{\lambda}{\gamma}\frac{F_1^2}{s} + \frac{\lambda+d}{\gamma}\frac{F_1}{s} - \frac{d}{\gamma}\frac{1-(1-x_2)s}{s}. \tag{S36}$$

Under the ansatz

$$F_1(s) = (\gamma/\lambda)sf'(s)/f(s), \tag{S37}$$

we express Eq. (S36) as

$$f''(s) + \frac{A}{s}f'(s) + \frac{B}{s^2}[1-(1-x_2)s]f(s) = 0, \tag{S38}$$

with

$$A = \frac{\gamma-(\lambda+d)}{\gamma}, \quad B = \frac{\lambda d}{\gamma^2}. \tag{S39}$$

Under the change in variable $z = 2\sqrt{B(1-x_2)s}$, Eq. (S38) reads

$$\frac{\mathrm{d}^2 f}{\mathrm{d}z^2} + \frac{2A-1}{z}\frac{\mathrm{d}f}{\mathrm{d}z} + \left(\frac{4B}{z^2} - 1\right) f = 0. \tag{S40}$$

Using now the ansatz

$$f = z^{1-A}\phi(z), \tag{S41}$$

and after some calculations, Eq. (S40) is recast into

$$z^2\phi''(z) + z\phi'(z) - (z^2 + \alpha^2)\phi(z) = 0, \tag{S42}$$

with $\alpha = (d-\lambda)/\gamma$. Note that under the stationarity condition $d > \lambda$, Eq. (2.3) of the main paper, $\alpha > 0$. Thus, the solution of Eq. (S42), for $\alpha$ not being an integer, is expressed in terms of the modified Bessel functions of the first kind $I_\alpha$ [7, Sec. 10.25][1]:

$$\phi(z) = I_\alpha(z) + D I_{-\alpha}(z). \tag{S43}$$

In Eq. (S43), $D$ is a constant that has to be defined from the initial condition $F_1(s=1) = x_1$. After some algebraic manipulations, we calculate constant $D$ to

$$D := D(x_1, x_2) = -\frac{[\alpha + 2\beta(1-x_1)] I_\alpha\left(2\sqrt{B(1-x_2)}\right) + 2\sqrt{B(1-x_2)} I'_\alpha\left(2\sqrt{B(1-x_2)}\right)}{[\alpha + 2\beta(1-x_1)] I_{-\alpha}\left(2\sqrt{B(1-x_2)}\right) + 2\sqrt{B(1-x_2)} I'_{-\alpha}\left(2\sqrt{B(1-x_2)}\right)}, \tag{S44}$$

with $\beta = \lambda/\gamma$.

Let us move now to the solution of the last equation (S32). Under the usual change of temporal variable $s = e^{-\gamma t}$, and by substituting ansatz (S37), Eq. (S32) is recast to

$$\frac{\partial_s F(s)}{F(s)} = \frac{rN_0}{\gamma}\frac{1}{s} - \frac{rN_0}{\lambda}\frac{f'(s)}{f(s)}, \quad F(s=1) = 1. \tag{S45}$$

The solution to Eq. (S45) is easily obtained:

$$\ln F(s) = \frac{rN_0}{\gamma}\ln s - \frac{rN_0}{\lambda}\ln\left(\frac{f(s)}{f(1)}\right). \tag{S46}$$

In solution (S46), we substitute function $f$ from Eq. (S41), resulting in

$$F(s) = s^{rN_0/\gamma}\left[s^{(1-A)/2}\frac{\phi\left(2\sqrt{B(1-x_2)s}\right)}{\phi\left(2\sqrt{B(1-x_2)}\right)}\right]^{-rN_0/\lambda}. \tag{S47}$$

After some calculations, the use of formula (S43) for $\phi\left(2\sqrt{B(1-x_2)s}\right)$, and the return to the original temporal variable $t$, we express Eq. (S47) equivalently as

$$F(x_1, x_2, t) = \left[\frac{\phi\left(2\sqrt{B(1-x_2)}\right) e^{(d-\lambda)t/2}}{I_\alpha\left(2\sqrt{B(1-x_2)}e^{-\gamma t/2}\right) + D(x_1,x_2) I_{-\alpha}\left(2\sqrt{B(1-x_2)}e^{-\gamma t/2}\right)}\right]^{rN_0/\lambda}. \tag{S48}$$

##### S5.3 Steady-state solution

For the steady-state solution, we have to calculate the long-time limit $t \to \infty$ of Eq. (S48). For this calculation, the following limiting form for modified Bessel functions is useful [7, Eq. 10.30.1]:

$$I_\alpha(z) \sim \left(\frac{1}{2}z\right)^\alpha \Big/ \Gamma(\alpha+1), \quad \text{for } z \to 0. \tag{S49}$$

[1] In case $\alpha$ is an integer, solution of Eq. (S42) should be expressed using the modified Bessel functions of both the first and second kind [8, Sec. 9.6]. However, for realistic estimates of parameter values, the case of integer $\alpha$ is unlikely, and thus we do not study it here.

Thus:

$$F(x_1,x_2)=\lim_{t\to\infty}F(x_1,x_2,t)=\lim_{t\to\infty}\left[\frac{\phi\left(2\sqrt{B(1-x_2)}\right)e^{(d-\lambda)t/2}}{\frac{(B(1-x_2))^{\alpha/2}e^{-(d-\lambda)t/2}}{\Gamma(1+\alpha)}+D\frac{(B(1-x_2))^{-\alpha/2}e^{(d-\lambda)t/2}}{\Gamma(1-\alpha)}}\right]^{rN_0/\lambda}=$$

$$=\left[\frac{\Gamma(1-\alpha)(B(1-x_2))^{\alpha/2}\phi\left(2\sqrt{B(1-x_2)}\right)}{D}\right]^{rN_0/\lambda},$$

and by using formula (S43) for $\phi\left(2\sqrt{B(1-x_2)}\right)$:

$$F(x_1,x_2)=\left[\Gamma(1-\alpha)(B(1-x_2))^{\alpha/2}\left(\frac{I_\alpha\left(2\sqrt{B(1-x_2)}\right)}{D(x_1,x_2)}+I_{-\alpha}\left(2\sqrt{B(1-x_2)}\right)\right)\right]^{rN_0/\lambda}. \tag{S50}$$

Last, by substituting $D(x_1,x_2)$ from Eq. (S44), steady-state joint PGF for TA and FD cell populations reads

$$F(x_1,x_2)=\left[h_{-\alpha}(x_2)\left(1-\frac{\alpha+2\beta(1-x_1)+q_{-\alpha}(x_2)}{\alpha+2\beta(1-x_1)+q_\alpha(x_2)}\right)\right]^{rN_0/\lambda}=$$

$$=\left(h_{-\alpha}(x_2)\frac{q_\alpha(x_2)-q_{-\alpha}(x_2)}{\alpha+2\beta(1-x_1)+q_\alpha(x_2)}\right)^{\frac{rN_0}{\lambda}}, \tag{S51}$$

with

$$h_{-\alpha}(x_2)=\Gamma(1-\alpha)(B(1-x_2))^{\alpha/2}I_{-\alpha}\left(2\sqrt{B(1-x_2)}\right), \tag{S52}$$

$$q_\alpha(x_2)=2\sqrt{B(1-x_2)}\frac{I'_\alpha\left(2\sqrt{B(1-x_2)}\right)}{I_\alpha\left(2\sqrt{B(1-x_2)}\right)}. \tag{S53}$$

Eq. (S51) constitutes our central result regarding the cell populations in a crypt that is in steady state.

**Useful formulas.** In this paragraph, we derive expressions for functions $h$, $q_\alpha$ and their derivatives that are useful in the calculation of limits of the following sections. By using the formula [9, entry 03.02.20.0005.01]

$$\frac{\mathrm{d}I_\alpha(z)}{\mathrm{d}z}=\frac{\alpha}{z}I_\alpha(z)+I_{\alpha+1}(z), \tag{S54}$$

Eq. (S53) is rewritten equivalently as

$$q_\alpha(x_2)=\alpha+2\sqrt{B(1-x_2)}\frac{I_{\alpha+1}\left(2\sqrt{B(1-x_2)}\right)}{I_\alpha\left(2\sqrt{B(1-x_2)}\right)}. \tag{S55}$$

By using Eq. (S54), as well as the alternative differentiation formula [9, entry 03.02.20.0004.01]

$$\frac{\mathrm{d}I_\alpha(z)}{\mathrm{d}z}=I_{\alpha-1}(z)-\frac{\alpha}{z}I_\alpha(z), \tag{S56}$$

we calculate the following derivatives

$$h'_{-\alpha}(x_2)=-\Gamma(1-\alpha)B(B(1-x_2))^{\frac{\alpha-1}{2}}I_{-\alpha+1}\left(2\sqrt{B(1-x_2)}\right), \tag{S57}$$

$$h''_{-\alpha}(x_2)=\Gamma(1-\alpha)B^2(B(1-x_2))^{\frac{\alpha-2}{2}}I_{-\alpha+2}\left(2\sqrt{B(1-x_2)}\right), \tag{S58}$$

$$q'_\alpha(x_2)=-2B+2B\frac{I_{\alpha+1}\left(2\sqrt{B(1-x_2)}\right)I_{\alpha-1}\left(2\sqrt{B(1-x_2)}\right)}{I^2_\alpha\left(2\sqrt{B(1-x_2)}\right)}, \tag{S59}$$

$$q''_\alpha(x_2) = -\frac{2B^2}{\sqrt{B(1-x_2)}}\left[\frac{I_{\alpha+2}\left(2\sqrt{B(1-x_2)}\right)I_{\alpha-1}\left(2\sqrt{B(1-x_2)}\right)}{I_\alpha^2\left(2\sqrt{B(1-x_2)}\right)}+\right.$$
$$\left.+\frac{I_\alpha\left(2\sqrt{B(1-x_2)}\right)I_{\alpha+1}\left(2\sqrt{B(1-x_2)}\right)}{I_\alpha^2\left(2\sqrt{B(1-x_2)}\right)} - \frac{2I_{\alpha+1}^2\left(2\sqrt{B(1-x_2)}\right)I_{\alpha-1}\left(2\sqrt{B(1-x_2)}\right)}{I_\alpha^3\left(2\sqrt{B(1-x_2)}\right)}\right]. \tag{S60}$$

##### S5.4 Validity check for the steady-state population of TA cells

By taking the limit $x_2 \to 1^-$, $F(x_1, x_2)$ of Eq. (S51) results in the steady-state PGF $F_{\mathrm{TA}}(x_1)$ for the TA cell population. By using the limiting form (S49) for modified Bessel functions and Eqs. (S52), (S55), we calculate

$$h_{-\alpha}(1^-) = 1, \quad q_\alpha(1^-) = \alpha, \quad q_{-\alpha}(1^-) = -\alpha, \tag{S61}$$

and thus

$$F_{\mathrm{TA}}(x_1) = \lim_{x_2\to 1^-} F(x_1,x_2) = \left(1 - \frac{\beta(1-x_1)}{\alpha+\beta(1-x_1)}\right)^{rN_0/\lambda} = \left(\frac{d-\lambda}{d-\lambda x_1}\right)^{rN_0/\lambda}. \tag{S62}$$

Eq. (S62) agrees with the exact steady-state solution for TA cell population (S14) derived independently in Sec. S3.

##### S5.5 Steady-state PGF for the FD cell population

By substituting $x_1 = 1$ in Eq. (S51), we obtain the steady-state PGF $F_{\mathrm{FD}}(x_2)$ for the FD cell population

$$F_{\mathrm{FD}}(x_2) = \left[h_{-\alpha}(x_2)\left(1 - \frac{\alpha + q_{-\alpha}(x_2)}{\alpha + q_\alpha(x_2)}\right)\right]^{rN_0/\lambda}. \tag{S63}$$

Using the limits of Eq. (S61), we see that $F_{\mathrm{FD}}(1^-) = 1$, which is an identity satisfied by all PGFs. From PGF (S63), we also calculate the average and variance of the FD cell population in steady-state, see [10, Sec. 4.4]:

$$F'_{\mathrm{FD}}(x_2) = \frac{rN_0}{\lambda}F_{\mathrm{FD}}(x_2)\left[\frac{h'_{-\alpha}(x_2)}{h_{-\alpha}(x_2)} - \frac{q'_{-\alpha}(x_2)}{q_\alpha(x_2)-q_{-\alpha}(x_2)} + \frac{q'_\alpha(x_2)}{q_\alpha(x_2)-q_{-\alpha}(x_2)}\frac{\alpha+q_{-\alpha}(x_2)}{\alpha+q_\alpha(x_2)}\right]. \tag{S64}$$

By using the limiting form (S49) and expressions Eqs. (S57), (S59), we calculate

$$h'_{-\alpha}(1^-) = \frac{B}{\alpha-1}, \quad q'_\alpha(1^-) = -\frac{2B}{\alpha+1}, \quad q'_{-\alpha}(1^-) = \frac{2B}{\alpha-1}. \tag{S65}$$

Thus:

$$\frac{h'_{-\alpha}(1^-)}{h_{-\alpha}(1^-)} = \frac{B}{\alpha-1}, \qquad \frac{\alpha+q_{-\alpha}(1^-)}{\alpha+q_\alpha(1^-)} = 0, \tag{S66}$$

$$\frac{q'_{-\alpha}(1^-)}{q_\alpha(1^-)-q_{-\alpha}(1^-)} = \frac{B}{\alpha(\alpha-1)}, \qquad \frac{q'_\alpha(1^-)}{q_\alpha(1^-)-q_{-\alpha}(1^-)} = -\frac{B}{\alpha(\alpha+1)}. \tag{S67}$$

By using Eqs. (S64), (S66) and (S67), we calculate the average number of FD cells in steady-state

$$m_{\mathrm{FD}} = F'_{\mathrm{FD}}(1^-) = \frac{rN_0}{\lambda}\left(\frac{B}{\alpha-1} - \frac{B}{\alpha(\alpha-1)}\right) = \frac{rN_0}{\lambda}\frac{B}{\alpha} = \frac{rdN_0}{\gamma(d-\lambda)}, \tag{S68}$$

which agrees with the result in Eq. (2.4) of the main paper. By differentiating expression (S64) again:

$$F''_{\mathrm{FD}}(x_2) = \frac{[F'_{\mathrm{FD}}(x_2)]^2}{F_{\mathrm{FD}}(x_2)} + \frac{rN_0}{\lambda}F_{\mathrm{FD}}(x_2)\left\{\frac{h''_{-\alpha}(x_2)}{h_{-\alpha}(x_2)} - \left[\frac{h'_{-\alpha}(x_2)}{h_{-\alpha}(x_2)}\right]^2 - \frac{q''_{-\alpha}(x_2)}{q_\alpha(x_2)-q_{-\alpha}(x_2)}+\right.$$
$$+\frac{q'_{-\alpha}(x_2)}{q_\alpha(x_2)-q_{-\alpha}(x_2)}\frac{q'_\alpha(x_2)}{q_\alpha(x_2)-q_{-\alpha}(x_2)} - \left[\frac{q'_{-\alpha}(x_2)}{q_\alpha(x_2)-q_{-\alpha}(x_2)}\right]^2+$$
$$+\left[\frac{q'_\alpha(x_2)}{q_\alpha(x_2)-q_{-\alpha}(x_2)}\right]'\frac{\alpha+q_{-\alpha}(x_2)}{\alpha+q_\alpha(x_2)}+$$
$$\left.+\frac{q'_\alpha(x_2)}{q_\alpha(x_2)-q_{-\alpha}(x_2)}\left[\frac{q'_{-\alpha}(x_2)}{\alpha+q_\alpha(x_2)} - \frac{q'_\alpha(x_2)}{\alpha+q_\alpha(x_2)}\frac{\alpha+q_{-\alpha}(x_2)}{\alpha+q_\alpha(x_2)}\right]\right\}. \tag{S69}$$

By using the limiting form (S49) and expressions Eqs. (S58), (S60), we calculate

$$h''_{-\alpha}(1^-) = \frac{B^2\Gamma(1-\alpha)}{\Gamma(3-\alpha)}, \quad q''_{\alpha}(1^-) = -\frac{4B^2}{(\alpha+1)^2(\alpha+2)}, \quad q''_{-\alpha}(1^-) = \frac{4B^2}{(\alpha-1)^2(\alpha-2)}. \tag{S70}$$

Thus, from Eqs. (S61), (S65), (S70), we obtain

$$\frac{h''_{-\alpha}(1^-)}{h_{-\alpha}(1^-)} = \frac{B^2}{(\alpha-1)(\alpha-2)}, \quad \frac{q''_{-\alpha}(1^-)}{q_{\alpha}(1^-)-q_{-\alpha}(1^-)} = \frac{2B^2}{\alpha(\alpha-1)^2(\alpha-2)}, \quad \frac{q'_{-\alpha}(1^-)}{\alpha+q_{\alpha}(1^-)} = \frac{B}{\alpha(\alpha-1)}. \tag{S71}$$

Using the derivative expressions (S64), (S69), and limit results (S66), (S67), (S71), we calculate the variance of FD cell population in steady-state

$$\sigma^2_{\mathrm{FD}} = F''_{\mathrm{FD}}(1^-) + F'_{\mathrm{FD}}(1^-) - \left[F'_{\mathrm{FD}}(1^-)\right]^2 = \frac{rN_0}{\lambda}\left[\frac{B}{\alpha} + \frac{B^2}{\alpha^2(\alpha+1)}\right]. \tag{S72}$$

Eq. (S72) is Eq. (2.9) of the main paper.

#### S5.6 Steady-state covariance of TA and FD cell populations

Since by Eq. (S51) we have determined the steady-state joint PGF $F(x_1, x_2)$ for TA and FD cell populations, we are able to calculate their steady-state covariance $\mathsf{Cov}(\mathrm{TA}, \mathrm{FD})$. First, we derive a general formula for the calculation of covariance from the joint probability-generating function. Definition relation for joint PGF $F(x_1, x_2)$ reads

$$F(x_1, x_2) = \sum_{n_1, n_2 \geq 0} x_1^{n_1} x_2^{n_2} P_{n_1, n_2}. \tag{S73}$$

By using their definition relation, we express the averages and the second joint moment of steady-state TA and FD cell populations in terms of PGF (S73) as

$$m_{\mathrm{TA}} = \sum_{n_1, n_2 \geq 0} n_1 P_{n_1, n_2} = \left.\frac{\partial F(x_1, x_2)}{\partial x_1}\right|_{x_1, x_2 = 1^-} \tag{S74}$$

$$m_{\mathrm{FD}} = \sum_{n_1, n_2 \geq 0} n_2 P_{n_1, n_2} = \left.\frac{\partial F(x_1, x_2)}{\partial x_2}\right|_{x_1, x_2 = 1^-} \tag{S75}$$

$$R_{\mathrm{TA,FD}} = \sum_{n_1, n_2 \geq 0} n_1 n_2 P_{n_1, n_2} = \left.\frac{\partial^2 F(x_1, x_2)}{\partial x_1 \partial x_2}\right|_{x_1, x_2 = 1^-} \tag{S76}$$

and for the covariance, which is the centered moment

$$\mathsf{Cov}(\mathrm{TA}, \mathrm{FD}) = R_{\mathrm{TA,FD}} - m_{\mathrm{TA}} m_{\mathrm{FD}} = \left.\frac{\partial^2 F(x_1, x_2)}{\partial x_1 \partial x_2}\right|_{x_1, x_2 = 1^-} - \left.\frac{\partial F(x_1, x_2)}{\partial x_1}\right|_{x_1, x_2 = 1^-} \left.\frac{\partial F(x_1, x_2)}{\partial x_2}\right|_{x_1, x_2 = 1^-} \tag{S77}$$

Since we have the explicit formula (S51), we determine it derivative with respect to $x_1$

$$\frac{\partial F(x_1, x_2)}{\partial x_1} = \frac{2\beta r N_0}{\lambda} \frac{F(x_1, x_2)}{\alpha + 2\beta(1-x_1) + q_{\alpha}(x_2)}, \tag{S78}$$

and by differentiating Eq. (S78) with respect to $x_2$

$$\frac{\partial^2 F(x_1, x_2)}{\partial x_1 \partial x_2} = \frac{2\beta r N_0}{\lambda} \frac{1}{\alpha + 2\beta(1-x_1) + q_{\alpha}(x_2)} \frac{\partial F(x_1, x_2)}{\partial x_2} - \frac{2\beta r N_0}{\lambda} \frac{F(x_1, x_2) q'_{\alpha}(x_2)}{\left[\alpha + 2\beta(1-x_1) + q_{\alpha}(x_2)\right]^2}. \tag{S79}$$

Since $F(1^-, 1^-) = 1$ (PGF property), $q_{\alpha}(1^-) = \alpha$ (Eq. (S61)), $q'_{\alpha}(1^-) = -2B/(\alpha+1)$ (Eq. (S65)), we calculate

$$\left.\frac{\partial F(x_1, x_2)}{\partial x_1}\right|_{x_1, x_2 = 1^-} = \frac{\beta r N_0}{\lambda \alpha} = \frac{r N_0}{d - \lambda}, \tag{S80}$$

and

$$\left.\frac{\partial^2 F(x_1,x_2)}{\partial x_1 \partial x_2}\right|_{x_1,x_2=1^-} = \frac{\beta r N_0}{\lambda\alpha} \left.\frac{\partial F(x_1,x_2)}{\partial x_2}\right|_{x_1,x_2=1^-} + \frac{\beta r N_0 B}{\lambda\alpha^2(\alpha+1)}. \tag{S81}$$

By substituting Eq. (S80) into Eq. (S81), we have

$$\left.\frac{\partial^2 F(x_1,x_2)}{\partial x_1 \partial x_2}\right|_{x_1,x_2=1^-} = \left.\frac{\partial F(x_1,x_2)}{\partial x_1}\right|_{x_1,x_2=1^-} \left.\frac{\partial F(x_1,x_2)}{\partial x_2}\right|_{x_1,x_2=1^-} + \frac{\beta r N_0 B}{\lambda\alpha^2(\alpha+1)}. \tag{S82}$$

Under Eq. (S82), Eq. (S77) reads

$$\mathsf{Cov}(\mathrm{TA},\mathrm{FD}) = \frac{\beta r N_0 B}{\lambda\alpha^2(\alpha+1)} = \frac{r\lambda d N_0}{\gamma(d-\lambda)^2(\alpha+1)}. \tag{S83}$$

Eq. (S83) is Eq. (2.10) of the main paper.

### S6 Steady-state probability distributions for cell populations

As in S5.1, we express the stationary joint PGF of stem, TA, and FD cell populations as

$$G(x_0,x_1,x_2) = x_0^{N_0} F(x_1,x_2), \tag{S84}$$

where $F(x_1,x_2)$ is the stationary joint PGF of TA and FD cells, given by Eq. (S51). From Eq. (S84), we deduce the PGF for total number of cells per crypt in steady-state:

$$F_{\mathrm{tot}}(x) = G(x,x,x) = x^{N_0} F(x,x), \tag{S85}$$

see also [6]. We recall that probability $P_n$ of a cell population being equal to $n$ is the coefficient of $x^n$ in the power series expansion of the corresponding PGF $F(x)$ [2, Sec. 2.2]:

$$F(x) = \sum_{n=0}^{\infty} P_n x^n. \tag{S86}$$

Thus, $P_n$ is determined by repeated differentiations of PGF $F(x)$, evaluated at $x=0$ [2, Sec. 2.5]:

$$P_n = \frac{F^{(n)}(0)}{n!}, \tag{S87}$$

where superscript $(n)$ denotes the $n$th order derivative of $F(x)$ with respect to its argument. A robust way of calculating (S87) is by Cauchy's integral formula [11, Th. 4.4]:

$$P_n = \frac{1}{2\pi i} \oint_C \frac{F(z)}{z^{n+1}} \mathrm{d}z, \tag{S88}$$

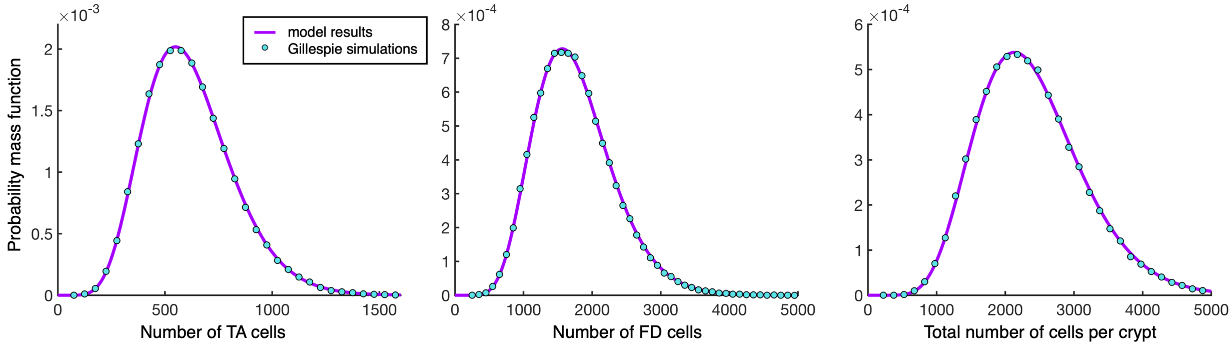


**Figure S1: Comparison between distributions from model solution and direct Gillespie simulations.** Steady-state probability mass function of TA, FD, and total cell populations in healthy human colonic crypts: $N_0 = 18$, $r = 1/2.5\mathrm{d}^{-1}$, $\lambda = 1/30\mathrm{h}^{-1}$, $\gamma = 1/3.5\mathrm{d}^{-1}$, $d$ is adjusted so that $N_{\mathrm{tot}} = 2392.1$. Number of Gillespie simulations performed 100,000. Simulation run time 500 days.

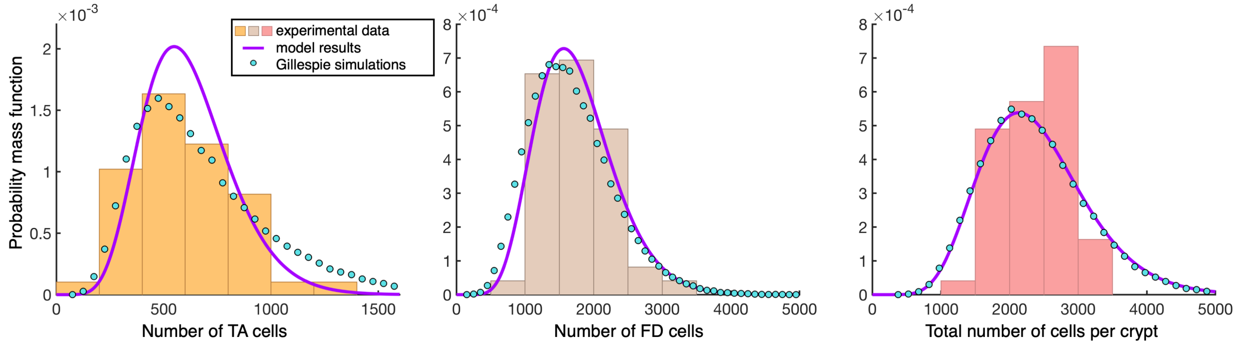


**Figure S2: Comparison between experimental data, model results, and Gillespie simulations with varying apoptosis rate.** Steady-state probability mass function of TA, FD, and total cell populations in healthy human colonic crypts: $N_0 = 18$, $r = 1/2.5\text{d}^{-1}$, $\lambda = 1/30\text{h}^{-1}$, $d$ is adjusted so that $N_{\text{tot}} = 2392.1$. For model results (purple curves) $1/\gamma = 3.5\text{d}$, while for Gillespie simulations (circles) $1/\gamma \sim \text{Uniform}[1\text{d}, 6\text{d}]$. Number of Gillespie simulations performed 100,000. Simulation run time 500 days. Histograms depict experimental data from Bravo and Axelrod [13]

where $i$ is the imaginary unit, and $C$ is a counterclockwise contour around the origin of the axes in complex plane, within the radius of convergence of $F(z)$. In order to calculate the probability mass functions (PMFs) of TA, FD or total number of cells per crypt in steady-state, we substitute the PGFs $F_{\text{TA}}(z)$ (Eq. (S14)), $F_{\text{FD}}(z)$ (Eq. (S63)), or $F_{\text{tot}}(z)$ (Eq. (S85)) in the right-hand side of Eq. (S88).

**Comparison to Gillespie simulations.** For the typical values of a human colonic crypt: $N_0 = 18$, $r = 1/2.5\text{d}^{-1}$, $\lambda = 1/30\text{h}^{-1}$, $\gamma = 1/3.5\text{d}^{-1}$, and $d$ adjusted so that the average total cell population per crypt is $N_{\text{tot}} = 2392.1$, we compare in Fig. S1 the steady-state PMFs of TA, FD and total cell populations as obtained from the model solution, to results obtained by 100,000 direct simulations of the branching process (S1) using Gillespie algorithm [12]. For TA, FD and total cell populations, the PMF is calculated by the numerical implementation of Cauchy's integral formula, Eq. (S88), over a circle contour around the origin of axes of unit radius.

### S7 Comparison of model results to experimental data

**Chi-square test between model results and data.** We perform a chi-square goodness of fit test [14, Sec.1.3.5.15], by using MATLAB function `chi2gof`, between the cell population distributions from model solution, for parameter values typical to healthy human colonic crypts, and experimental data [13, 15], shown in Fig. 3 of the main paper. The calculated $p$-values from the test are 0.24, 0.67 and 0.46, for TA,

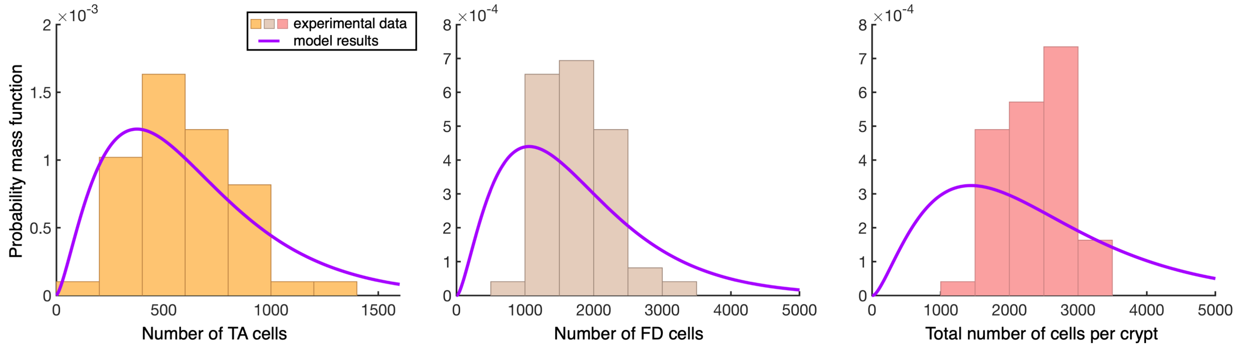


**Figure S3: Comparison between experimental data and distributions from model solution for 5 stem cells.** Steady-state probability mass function of TA, FD, and total cell populations in healthy human colonic crypts: $N_0 = 5$, $r = 1/2.5\text{d}^{-1}$, $\lambda = 1/30\text{h}^{-1}$, $\gamma = 1/3.5\text{d}^{-1}$, $d$ is adjusted so that $N_{\text{tot}} = 2392.1$. Histograms depict experimental data from Bravo and Axelrod [13]

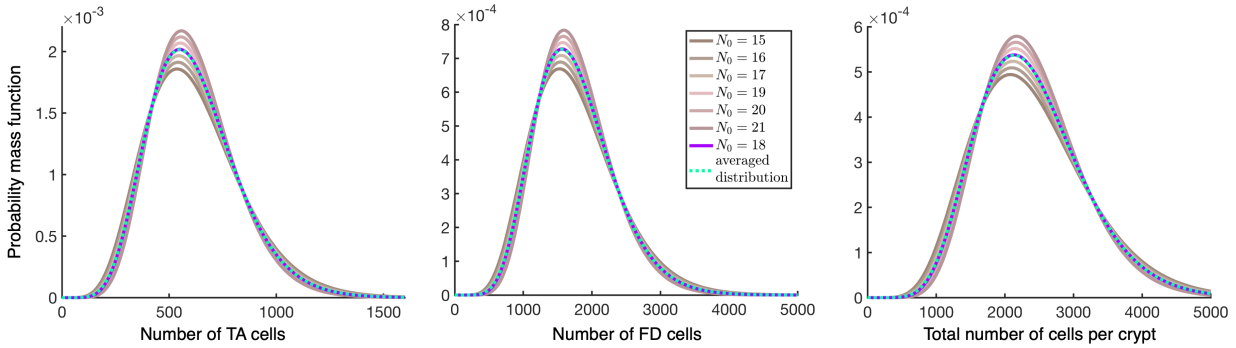


**Figure S4: Distributions from model solution for different number of stem cells, and their variances.** Steady-state probability mass function of TA, FD, and total cell populations in healthy human colonic crypts: $r = 1/2.5\mathrm{d}^{-1}$, $\lambda = 1/30\mathrm{h}^{-1}$, $\gamma = 1/3.5\mathrm{d}^{-1}$, $d$ is adjusted so that $N_{\mathrm{tot}} = 2392.1$. Solid lines depict distributions with $N_0 = 15 - 21$. Dashed line depicts the average of distributions with $N_0 = 15 - 21$.

FD and total cell populations respectively. Thus, at 5% significance level, chi-square test does not reject the hypotheses that experimental data for TA, FD and total cell populations come from the respective distributions predicted by our model.

**Model results for varying $\gamma$.** We perform 100,000 direct Gillespie simulations of branching process (S1), allowing apoptosis rate $\gamma$ to vary from crypt to crypt, with $1/\gamma \sim \mathrm{Uniform}[1\mathrm{d}, 6\mathrm{d}]$. The resulting probability distributions are shown in Fig. S2, along with our model results for $1/\gamma = 3.5\mathrm{d}$ and experimental data [13]. We observe that the distributions from simulations with varying $\gamma$ stay close to data, while the distribution for total cell population effectively coincides with the corresponding distribution for constant $\gamma$.

**Model results for $N_0 = 5$.** There is an ambiguity regarding the number of stem cells per crypt, since lower values for $N_0$ around 5 cells have also been reported. In Fig. S3, we show the calculated distributions for $N_0 = 5$ along with the experimental data, and we observe that they are not in agreement. This is corroborated by the chi-square goodness of fit test: the $p$-values calculated by MATLAB function `chi2gof` are 0.097, 0.003 and 0.028 for TA, FD and total cell populations respectively. Thus, at a 5% significance level, chi-square test disproves the hypothesis that experimental data for FD and total cell populations come from the respective distributions predicted by our model. The hypothesis is not rejected only for the TA cell population. Furthermore, the model for $N_0 = 5$ predicts coefficients of variation for all three cell populations $\sim 63\%$. This is significantly higher than the values of coefficients of variation calculated from experimental data (see Table 1 of the main paper).

**Model results for varying $N_0$.** We examine the stability of our results under variations in stem cell number $N_0$ from crypt to crypt. In Fig. S4, we show the cell population distributions determined from the model solution for $N_0 \in [15, 21]$, and corresponding averaged distributions. We see that the averaged distributions coincide with the distributions for $N_0 = 18$.

### References


[1] Allen, L. J. S. *An Introduction to Stochastic Processes with Applications to Biology* (CRC Press, Boca Raton, 2010), 2nd edn.

[2] Bailey, N. T. J. *The elements of Stochastic Processes with applications to the natural sciences* (John Wiley and sons, New York, London, Sydney, 1964).

[3] Polyanin, A. D., Zaitsev, V. F. & Moussiaux, A. *Handbook of First Order Partial Differential Equations* (Taylor and Francis, New York, 2002).

[4] Avanzini, S. *et al.* A mathematical model of ctDNA shedding predicts tumor detection size. *Sci. Adv* **6**, eabc4308 (2020).

[5] Antal, T. & Krapivsky, P. L. Exact solution of a two-type branching process: Models of tumor progression. *Journal of Statistical Mechanics: Theory and Experiment* **2011**, P08018 (2011).

[6] Antal, T. & Krapivsky, P. L. Exact solution of a two-type branching process: Clone size distribution in cell division kinetics. *Journal of Statistical Mechanics: Theory and Experiment* **2010**, P07028 (2010).

[7] Olver, F. W. J. *et al.* NIST Digital Library of Mathematical Functions. http://dlmf.nist.gov/, Release 1.1.7 of 2022-10-15 (2022).

[8] Abramowitz, M. & Stegun, I. A. *Handbook of Mathematical Functions with Formulas, Graphs, and Mathematical Tables* (Dover Publications (1983 reprint), 1964), 10th edn.

[9] Wolfam Research Inc. The mathematical functions site (2023). URL http://functions.wolfram.com/.

[10] Fewster, R. *Stochastic Processes (course notes STATS325)* (Department of Statistics, University of Auckland, Auckland, NZ, 2014).

[11] Dorin, A. & Bagdasar, O. *Recurrent Sequences* (Springer, Cham, Switzerland, 2020).

[12] Nezar. Gillespie Stochastic Simulation Algorithm (https://github.com/nvictus/Gillespie) (2023).

[13] Bravo, R. & Axelrod, D. E. A calibrated agent-based computer model of stochastic cell dynamics in normal human colon crypts useful for in silico experiments. *Theoretical Biology and Medical Modelling* **10**, 66 (2013).

[14] NIST/SEMATECH. e-Handbook of Statistical Methods (2023). URL http://www.itl.nist.gov/div898/handbook/.

[15] Yang, J., Axelrod, D. E. & Komarova, N. L. Determining the control networks regulating stem cell lineages in colonic crypts. *Journal of Theoretical Biology* **429**, 190–203 (2017).
